# Single-cell gene networks nominate *IKZF1* as an Alzheimer’s microglial regulator

**DOI:** 10.64898/2026.07.14.738463

**Authors:** Çağrı Özkurt

## Abstract

**Background:** Microglia drive neuroinflammation in Alzheimer’s disease (AD), yet no approved therapy targets this compartment. Human genome-wide association studies consistently implicate innate immune loci in AD risk, establishing microglial transcriptional programs as therapeutically relevant but pharmacologically underexploited targets.

**Objective:** We sought to identify transcription factors (TFs) governing microglial state transitions computationally and to nominate structurally tractable drug repurposing candidates.

**Methods:** We applied trajectory inference (PAGA), pseudobulk DESeq2, pySCENIC gene regulatory network (GRN) inference, CellChat, and virtual screening of 1,962 approved compounds to 236,002 microglial nuclei from 84 donors (SEA-AD atlas).

**Results:** *IKZF1* was the sole target TF retained under cisTarget v10 motif constraints, with peak regulon activity in LateAD-DAM (pseudotime ρ = +0.309) and replication in an independent bulk cohort (GSE95587; adjusted P value =.004). CellChat identified SLIT2→ROBO2 from multiple neuron subtypes (predominantly inhibitory interneurons) as the top predicted pathway to microglia. Tafamidis (→IRF8) and diflunisal (→PPARG) were top virtual screening hits; all evaluated compounds failed the pre-specified selectivity threshold.

**Conclusions:** *IKZF1* is prioritised as a candidate late-disease microglial TF, supported by six convergent evidence dimensions including independent bulk replication. Tafamidis and diflunisal are low-confidence repurposing hypotheses requiring experimental validation.

## Introduction

Alzheimer’s disease (AD) is the leading cause of dementia, affecting tens of millions of people worldwide with prevalence expected to increase substantially as global populations age. Despite approval of amyloid-targeting immunotherapies, disease-modifying options remain limited and do not engage the neuroinflammatory component of disease [1]. Human genome-wide association studies have consistently implicated innate immune loci — including *TREM2*, *CLU*, *BIN1*, and *INPP5D* — in AD risk [2,3], establishing the microglial compartment as a therapeutically relevant but pharmacologically underexploited target.

Microglia, the brain’s resident immune sentinels, undergo complex state transitions in Alzheimer’s disease. They shift from homeostatic surveillance to a spectrum of disease-associated states, including interferon-responsive (IRM), disease-associated (DAM), lipid-associated (LAM), and terminal late-disease phenotypes [2,4]. Identifying the transcription factors (TFs) governing these transitions is critical for understanding — and potentially reversing — pathological microglial activation.

Due to significant species-specific divergence in microglial signatures [3], we focus exclusively on human-derived single-nucleus data. Master regulators like *SPI1*/*PU.1*, *IRF8*, and *PPARG* have been implicated in various microglial states [3,5], but prioritizing these candidates for pharmacological intervention remains a significant challenge. These TFs possess structurally defined DNA-binding domains that are in principle targetable by small molecules [6,7].

Gene regulatory network (GRN) inference offers a principled route to TF prioritisation by identifying TFs that are causally active rather than merely differentially expressed [8]. Combined with structural druggability assessment and virtual screening, GRN analysis can narrow the hypothesis space from hundreds of candidate regulators to a handful of mechanistically credible, structurally tractable targets [9].

Here we apply the DAM-DRUG framework to the SEA-AD multi-regional atlas [10], the largest publicly available single-nucleus microglial dataset. The novelty of this work lies in the multi-modality convergence logic that links established tools—pySCENIC, CellChat, AutoDock Vina, and CellOracle—into a ranked evidence hierarchy. This framework requires a candidate TF to be supported across trajectory, regulon, chromatin, bulk replication, and perturbation evidence before advancing to structural screening.

## Methods

All analyses were executed in Apptainer containers on the TRUBA ARF HPC cluster (TÜBİTAK ULAKBİM) or a local macOS workstation. Container images, exact software version pins, and step-by-step execution instructions are documented in the GitHub repository README.

### Dataset acquisition and preprocessing

Single-nucleus RNA-seq data were obtained from the SEA-AD multi-regional microglial release ([10]; AWS S3 bucket s3://sea-ad-single-cell-profiling; 236,002 nuclei, 84 donors, 10 regions). MTG paired multiome RNA and ATAC objects were downloaded from the same portal. Pre-annotated Supertype labels were mapped to six functional substates by direct lookup (**Supplementary Table S1**). Per-dataset QC and batch correction were not repeated (Allen Institute harmonised pipeline; Scrublet + scVI). Disease group classification uses Cognitive Status (Dementia/No dementia) for cohort description, Overall ADNC for neuropathological severity, and Severely Affected Donor (High ADNC + severe neurodegeneration) for ATAC DA analysis. The cohort includes donors of both sexes (female: 27/42 Dementia, 24/42 No dementia; p = 0.503); sex was retained as a covariate (sex_bin) in all differential expression and ATAC models. Donor inclusion criteria per analysis are in **Supplementary Table S2**.

### Trajectory inference

PAGA [11] was applied to the 10-dimensional scVI embedding. Diffusion pseudotime was rooted at the Homeostatic nucleus with maximal *P2RY12* expression, utilizing its established role as the canonical marker for the ramified, surveillant state [12,13]. This selection aligns with histological evidence in AD showing that microglial P2RY12 expression is high in homeostatic environments and progressively lost during pathological activation and plaque association [14,15].

### Differential gene expression

Pseudobulk DESeq2 (pydeseq2) contrasted DAM, IRM, and LAM vs Homeostatic using design ∼group + sex_bin + age_z; pseudobulk samples summed raw UMI counts per donor across all brain regions. LateAD-DAM was excluded (median 4 cells/donor). DAM-IRM was excluded as a primary contrast due to its hybrid co-expression profile. Minimum 10 cells/donor per state required for inclusion. Bulk replication: GSE95587 (fusiform gyrus; n=117; design: ∼condition + sex_bin + age_z + rin_z); acceptance criterion adjusted P value <.05, log₂FC > 0.5.

### Gene regulatory network inference

pySCENIC v0.12.1 (5 GRNBoost2 seeds; consensus edges in ≥3/5 runs; 1,458,113 edges; 100,000-cell random subsample from 236,002; RcisTarget against cisTarget v10/HOCOMOCO v11; 80% gene-mapping threshold; AUCell top 5% per cell). 46 regulons passed pruning. Regulon-pseudotime Spearman correlations BH-corrected across 46 tests. BHLHE40/41 rescue: pseudobulk Spearman co-expression across 84 donors; top 200 co-expressed genes tested by Fisher’s exact vs all expressed genes.

### ATAC chromatin accessibility validation

Microglia isolated from MTG ATACseq h5ad; pseudobulked per donor (84 of 86 donors (2 excluded: no Severely Affected label)); Severely Affected (n=11) vs non-Severely-Affected (n=73). Mann-Whitney U (two-sided) per peak across 218,882 peaks; BH FDR correction; DAPs: FDR < 0.05, |log₂FC| > 1 (1,191 total; 56 AD-upregulated). FIMO (MEME suite; HOCOMOCO v11 motifs; P value < 1×10⁻⁴) on 56 AD-upregulated vs 5,000 FDR > 0.2 background peaks; enrichment = hit-rate ratio (hit-rate ratio descriptive).

### Cell–cell communication analysis

CellChat v2 [16]; R 4.3; 45,000 MTG nuclei (up to 5,000 per cell type; 15 types); Secreted Signaling subset (1,280 L-R pairs); computeCommunProb with triMean; interactions ranked by cumulative probability (no multiple-testing correction).

### Protein structure preparation, druggability, and virtual screening

Protein structures were obtained from the AlphaFold Protein Structure Database [17], utilizing the human proteome v6 release generated by the AlphaFold2 algorithm [18]. Structures were trimmed to functional domains using a confidence threshold of pLDDT ≥ 50 to exclude highly disordered regions. Models were prepared using PDBFixer [19] to add missing atoms and determine terminal ionization states at pH 7.4. Structures underwent implicit-solvent (GBn2) energy minimization (200 steps) using OpenMM 8.1 [20] with the AMBER14 force field. fpocket 4.0 was employed to identify potential binding sites, with druggability scores parsed per-pocket. Ligand libraries assembled from ChEMBL v34: Tier-1 (285 CNS-approved, ATC-N, Ro5); Tier-2 (1,677; Ro5 + CNS MPO proxy ≥ 3 [21]; IRF8/IKZF1 only). 3D conformers via Open Babel 3.1 (--gen3d-p 7.4); PDBQT via Meeko 0.5 [22]. AutoDock Vina 1.2 (--exhaustiveness 32 -- num_modes 10); GNINA 1.0 CNN rescoring (crossdock_default2018; top 30 poses). Consensus score S = 0.4×(|Vina|/10) + 0.6×CNN. MM-GBSA: gmx_MMPBSA 1.6.4 [23] (GROMACS 2024.1; AMBER99SB-ILDN + GAFF2; 1 ns NPT; igb=2; 250 frames). Selectivity: SI = |ΔG_on-target|/|ΔG_off-target| vs DRD2/6CM4, HTR2A/6A94, hERG/7CN1 [24]; advancement criterion SI > 2.0 (none met). 100 ns MD stability: AMBER99SB-ILDN + GAFF2 + TIP3P; Ewald PME; 2 fs timestep; core RMSD on 10% lowest-RMSF residues. Grid box coordinates and pocket parameters are in **Supplementary Table S3**; pre-specified thresholds and consensus scoring metrics are in **Supplementary Table S4**.

### CellOracle in-silico TF perturbation

In-silico transcription factor (TF) knockouts (KO) were performed using CellOracle v0.12 [25]; on a 20,000-cell stratified subsample of the microglial atlas. The base gene regulatory network (GRN) was constructed from the 5-seed pySCENIC consensus adjacency, restricted to the top 500 targets per TF based on importance scores. Simulated KO was executed by setting the expression of the target TF to zero, with signal propagation depth set to n_propagation_=3.

The magnitude of the resulting state transition was quantified as the magnitude of the mean displacement vector per substate in the UMAP embedding (mean |ΔUMAP|). It is important to note that because UMAP is a non-linear manifold learning technique that prioritizes local connectivity over global distance preservation [26], distances in this embedding space are often distorted and do not faithfully represent biological distances [27]. Consequently, these perturbation metrics are utilized exclusively for the ordinal ranking of TF candidates within specific substates rather than as linear quantitative measurements of biological effect size.

### Statistical software

Python 3.10; scanpy 1.11.5, pydeseq2 0.5.4, pyscenic 0.12.1, CellChat 2.2.0 via R 4.3.3. All stochastic steps used fixed seeds (**Supplementary Table S5**).

## Results

### Single-nucleus atlas and microglial substate annotation

We obtained 236,002 microglial nuclei from 84 postmortem donors spanning Braak stages 0–VI (42 Dementia, 42 No dementia; age and PMI balanced between groups; **Table 1**). Pre-annotated SEA-AD Supertype labels were mapped to six functional substates [10](**Supplementary Table S1**): Homeostatic (HM; n=7,909), Disease-associated microglia (DAM; n=141,248) [2], DAM-IRM (n=53,815), Interferon-responsive microglia (IRM; n=29,805), LateAD-DAM (n=1,458), and Lipid-associated microglia (LAM; n=1,767). DAM-IRM, a hybrid state co-expressing DAM (*TREM2*, *APOE*) and IRM (*IFIT1*, *MX1*) markers, was retained in AUCell, pseudotime, and CellOracle analyses but excluded from primary DGE contrasts. LateAD-DAM and LAM were present in 66/84 and 68/84 donors respectively (median 4 and 5 cells/donor), constraining donor-level pseudobulk analyses. LDAM module scores [28] confirmed substate identity (LAM = 1.13 >> IRM = 0.59 > DAM = 0.08 > HM = −0.11; **Figure 1A-B; Supplementary Fig. S1**). Substate cell counts by region are in **Supplementary Table S6**.

**Figure 1.**
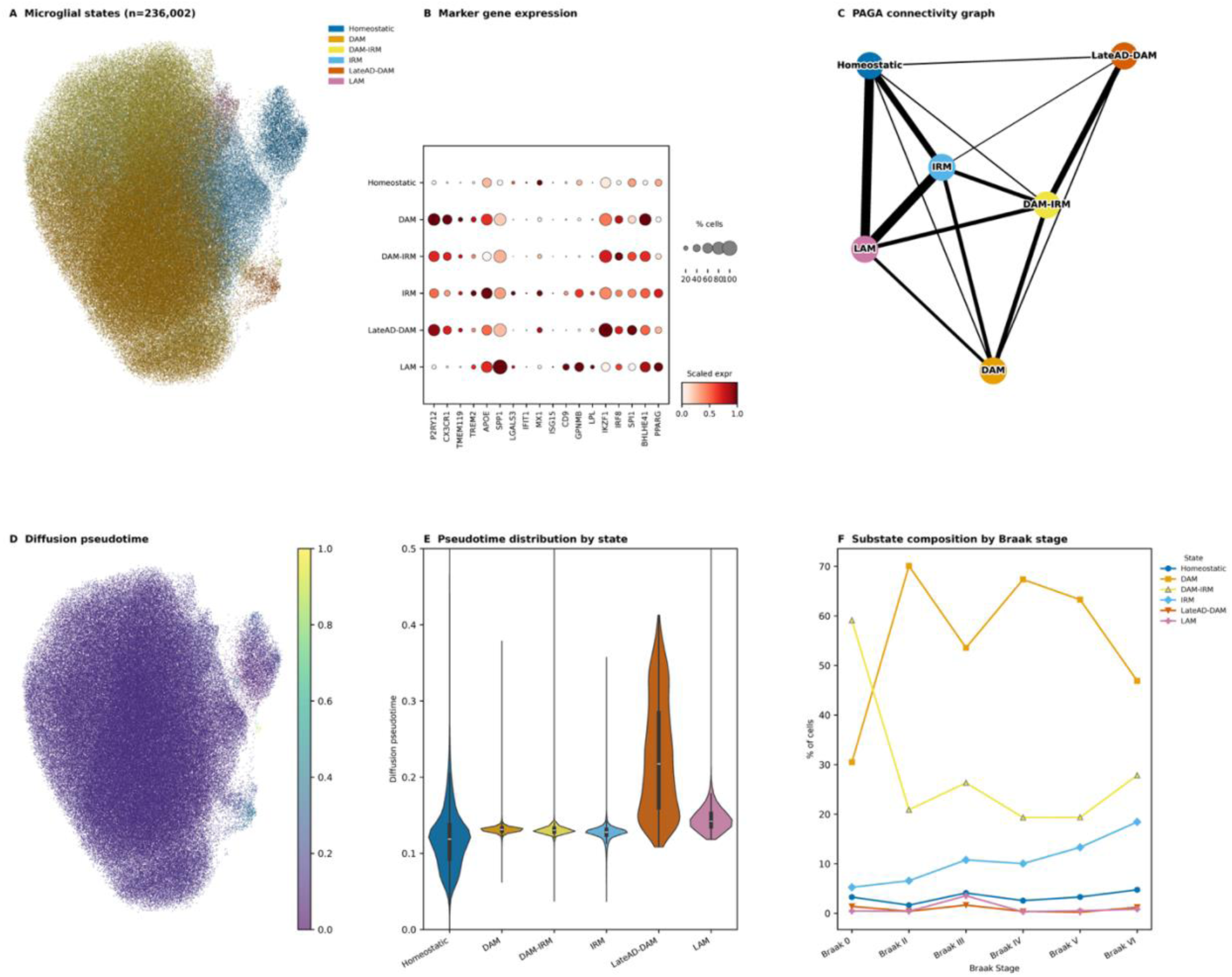
Single-nucleus microglial atlas and substate annotation. **(A)** UMAP of 236,002 microglial nuclei (84 donors, SEA-AD atlas) coloured by substate. **(B)** Dot plot of canonical marker gene expression; dot size = % cells expressing; colour = mean scaled expression. **(C)** PAGA connectivity graph; edge weight proportional to substate connectivity. **(D)** Diffusion pseudotime (DPT) overlaid on UMAP; LateAD-DAM occupies the highest values (median = 0.218), confirming it as the terminal state. **(E)** Violin plots of DPT per substate (ordering: HM→IRM→DAM≈DAM-IRM→LAM→LateAD-DAM). **(F)** Substate composition (%) across Braak stages 0–VI.

**Table 1.** Demographic and Sample Distribution of the SEA-AD Microglial Dataset. Summary of microglial nuclei (n = 236,002) obtained from 84 postmortem donors across the Alzheimer’s disease (AD) neuropathological continuum. Donors are stratified by clinical Cognitive Status (42 “Dementia” cases vs. 42 “No dementia” controls). Cell counts are provided for the six functional substates defined by mapping SEA-AD Supertypes: Homeostatic (HM), Disease-associated microglia (DAM), DAM-IRM, Interferon-responsive microglia (IRM), Late-stage DAM (LateAD-DAM), and Lipid-associated microglia (LAM). While all 84 donors contributed cells to the primary states (HM, DAM, IRM), the terminal states (LAM and LateAD-DAM) exhibit characteristic sparsity across the cohort. Full regional distributions and donor-level metadata are provided in Supplementary Data.

| State | n Donors | Mean |  |  |  |
| --- | --- | --- | --- | --- | --- |
|  |  | Cells | Median Cells | Min Cells | Max Cells |
| **Homeostatic** | 84 | 94.15 | 63.0 | 12 | 389 |
| **DAM** | 84 | 1,681.52 | 1,503.0 | 119 | 4,981 |
| **DAM-IRM** | 84 | 640.65 | 388.5 | 71 | 2,394 |
| **IRM** | 84 | 354.82 | 272.5 | 48 | 1,366 |
| **LateAD-DAM** | 84 | 17.36 | 4.0 | 0 | 221 |
| **LAM** | 84 | 21.04 | 5.0 | 0 | 608 |

### Trajectory inference defines a linear disease-progression axis

PAGA and diffusion pseudotime (DPT) defined the ordering HM → IRM → DAM-IRM ≈ DAM → LAM → LateAD-DAM (DPT = 0.119–0.218), with the largest pseudotime gap at the LAM→LateAD-DAM transition (Δ = 0.076), confirming LateAD-DAM as the terminal state (**Figure 1C-E**). PAGA connectivity analysis revealed LAM as the central transcriptional hub of the microglial state graph, with maximum connectivity scores to both HM (1.00) and IRM (1.00), reflecting substantial transcriptional overlap between these states. DAM was connected to the graph primarily through DAM-IRM (connectivity = 0.49), which itself bridged IRM (0.43) and LateAD-DAM (0.65), consistent with DAM-IRM occupying an intermediate position between the IRM and disease-terminal lineages. Because PAGA encodes undirected transcriptional proximity rather than directed state transitions, the disease-progression axis is defined by DPT ordering rather than PAGA topology.

### Pseudobulk differential expression defines state-specific TF signatures

Pseudobulk DESeq2 analysis (DAM, IRM, and LAM vs Homeostatic; 84 donors as replicates; LateAD-DAM excluded due to sparse per-donor counts; DAM-IRM excluded as a primary contrast owing to its hybrid profile) identified state-specific TF signatures (top markers in **Table 2**). Among the nine target TFs: *BHLHE41* showed the strongest pan-disease signal (DAM log₂FC = +2.55, adjusted P value = 6.9 × 10^-195^; IRM +1.89; LAM +2.44); *IRF8* was upregulated in all three states (DAM +1.70, adjusted P value = 8.5 × 10^-94^); *PPARG* showed a non-monotonic pattern (DAM −0.82; LAM +1.15), its established repression during acute neuroinflammation and its critical re-activation during the induction of lipid-processing and immunometabolic reprogramming in the LAM state [28,29]; *IKZF1* was DAM/IRM-specific (DAM +0.75, adjusted P value = 6.9 × 10^-51^; IRM +0.61; absent from LAM), consistent across all 84 donors.

**Table 2.** Top differentially expressed genes and prioritized transcription factors across microglial substates. The table summarizes the results of pseudobulk DESeq2 analysis for the three primary disease-state contrasts (DAM, IRM, and LAM) versus the Homeostatic reference. Genes are ranked by log_2_ fold-change (log_2_FC) within each contrast. All reported genes reached statistical significance at p_adj_<0.05. Differential expression was calculated across 84 donors (for DAM and IRM) and 27 donors (for LAM), treating each donor as a biological replicate to account for inter-individual variability. Functional annotations highlight the shift from homeostatic surveillance to states characterized by extracellular matrix remodeling (ADAMTSL2), synaptic modulation (SYNDIG1), and lipid-associated inflammation (CCL3L1). Notably, the human-specific upregulation of CX3CR1 is observed in the DAM state, diverging from mouse models where it is typically downregulated during disease progression. LateAD-DAM was excluded from the pseudobulk contrast due to high sparsity across the donor cohort. Only protein-coding genes are shown; lncRNAs and pseudogenes are excluded. Full results (6,712 / 3,956 / 2,985 upregulated significant genes for DAM / IRM / LAM contrasts respectively) are available at Zenodo (https://zenodo.org/records/19559648). Design formula: ‘∼group + sex_bin + age_z’; BH FDR correction applied genome-wide per contrast.

| Contrast | Gene | log<sub>2</sub>FC | p<sub>adj</sub> | Function |
| --- | --- | --- | --- | --- |
| **DAM** | ADAMTS | +4.37 | 6.6 × | Extracellular matrix |
|  | L2 |  | 10<sup>-146</sup> | glycoprotein; TGFβ signalling modulator |
| **DAM** | SYNDIG1 | +4.33 | 2.3 × | Synapse development; |
|  |  |  | 10<sup>-255</sup> | transmembrane protein |
| **DAM** | CX3CR1 | +4.20 | 5.9 × | Fractalkine receptor; human |
|  |  |  | 10<sup>-182</sup> | DAM activation marker |
| **DAM** | MMRN1 | +4.16 | 6.2 × 10<sup>-70</sup> | Multimerin-1; extracellular matrix/platelet protein |
| **DAM** | KCNIP1 | +4.13 | 2.8 × | Potassium channel interacting |
|  |  |  | 10<sup>-217</sup> | protein; Kv4 channel regulator |
| **IRM** | SYNDIG1 | +3.75 | 7.5 × | Synapse development; |
|  |  |  | 10<sup>-149</sup> | transmembrane protein |
| **IRM** | ADAMTS | +3.72 | 3.8 × 10<sup>-94</sup> | Extracellular matrix glycoprotein |
|  | L2 |  |  |  |
| IRM | CLEC19A | +3.67 | $2.2 \times 10^{-44}$ | C-type lectin domain family member; immune receptor |
| IRM | CYP19A1 | +3.66 | $2.5 \times 10^{-42}$ | Aromatase; oestrogen biosynthesis enzyme |
| IRM | MORC1 | +3.65 | $9.1 \times 10^{-67}$ | GHKL ATPase; germline/epigenetic regulator |
| LAM | RGCC | +5.00 | $1.5 \times 10^{-56}$ | Regulator of cell cycle; apoptosis modulator |
| LAM | CCL3L1 | +4.82 | $1.5 \times 10^{-16}$ | CC chemokine; CCR1/CCR5 ligand; pro-inflammatory |
| LAM | EGR3 | +4.74 | $4.9 \times 10^{-29}$ | Early growth response TF; lymphocyte development |
| LAM | FLT1 | +4.67 | $3.3 \times 10^{-35}$ | VEGF receptor 1; angiogenesis/inflammatory signalling |
| LAM | CCL4L2 | +4.66 | $4.9 \times 10^{-11}$ | CC chemokine; CCR5 ligand; macrophage inflammatory protein |

### Gene regulatory network inference identifies IKZF1 as the primary disease regulon

pySCENIC (5-seed GRNBoost2 consensus; 1.46 M co-expression edges; RcisTarget against cisTarget v10/HOCOMOCO v11; **Supplementary Methods SM1**) identified 46 regulons above the 80% gene-mapping threshold. Of the nine target TFs, only *IKZF1* was retained; the remaining eight failed motif-database coverage—a bottleneck of cisTarget v10 annotation, not GRNBoost2 stochasticity (confirmed across 5 seeds; **Supplementary Methods SM2**; **Supplementary Table S7**). IKZF1 prioritisation is therefore conditional on this database and does not imply other TFs lack regulatory roles. A motif-agnostic pseudobulk co-expression rescue analysis for *BHLHE40*/*41* found no enrichment of DAM/LAM markers in either TF’s top co-expressed genes (Fisher’s exact OR = 0.00, P value = 1.0), consistent with these TFs acting within cell-state transitions rather than as donor-level axes (**Supplementary Figs. S2, S3**). The *IKZF1*(+) regulon peaked in LateAD-DAM (AUC = 0.153) and showed the only positive pseudotime correlation among the 46 retained regulons (Spearman ρ = +0.309; anti-conservative CI due to within-donor clustering; **Figure 2; Supplementary Fig. S4**).

**Figure 2.**
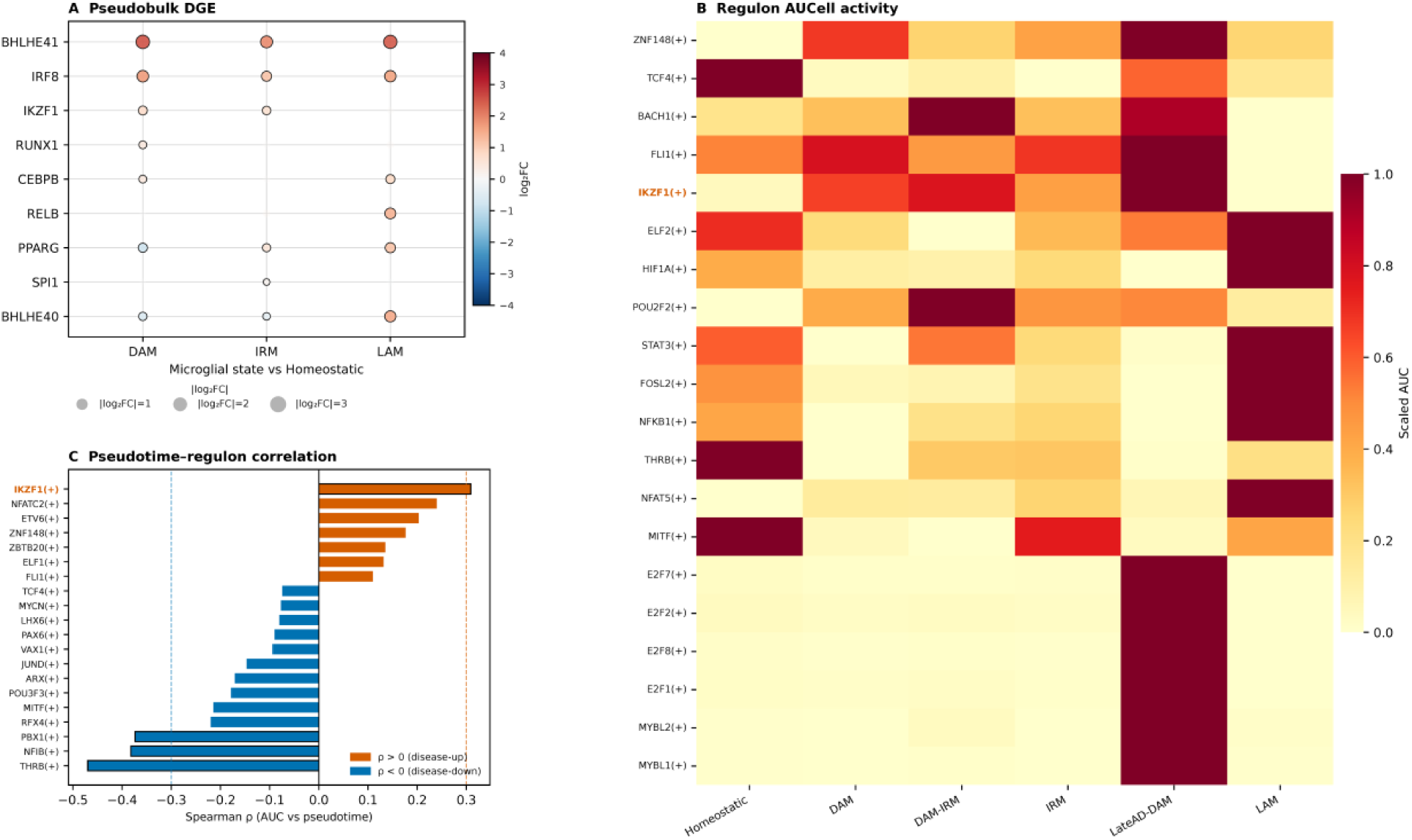
Multi-modality transcription factor prioritisation across microglial substates. **(A)** Pseudobulk DESeq2 results for nine target TFs (DAM, IRM, LAM vs Homeostatic); dot size = −log10(adjusted P value), colour = log₂FC. **(B)** AUCell regulon activity heatmap for 46 pySCENIC regulons across six substates. **(C)** Spearman correlation of regulon AUCell scores with DPT (BH-corrected); IKZF1(+) is the only target TF regulon with a significant positive correlation (ρ = +0.309).

### Multi-evidence prioritisation of IKZF1

Six analyses converge on *IKZF1* (**Figure 3D**; **Supplementary Fig. S5**), classified explicitly by epistemic status. Four internal SEA-AD RNA analyses (exploratory; non-independent; same 84 donors): pseudobulk DGE upregulation in DAM and IRM; *IKZF1*(+) regulon peaking in LateAD-DAM; monotonic pseudotime correlation (ρ = +0.309); and CellOracle KO producing the largest LateAD-DAM UMAP displacement among three TFs tested (|ΔUMAP| = 0.047 vs *IRF8* = 0.026 vs *SPI1* = 0.013; ordinal ranking only). Orthogonal corroboration from SEA-AD ATAC (same donors, different assay): *IKZF1* motif 3.90× enriched in AD-upregulated open chromatin peaks (descriptive; no GC-matched background; **Supplementary Methods SM3**). The only external replication: bulk RNA-seq upregulation in AD fusiform gyrus (GSE95587; n=117 independent donors; log₂FC = +0.638, adjusted P value =.004). Literature triangulation: IKZF1 protein elevation in AD hippocampus confirmed by western blot [30]. Analyses 1–5 are non-independent; item 6 is the sole external replication. *IRF8*, *SPI1*, and *BHLHE41* also reached significance in the independent bulk cohort, confirming the broader TF programme.

**Figure 3.**
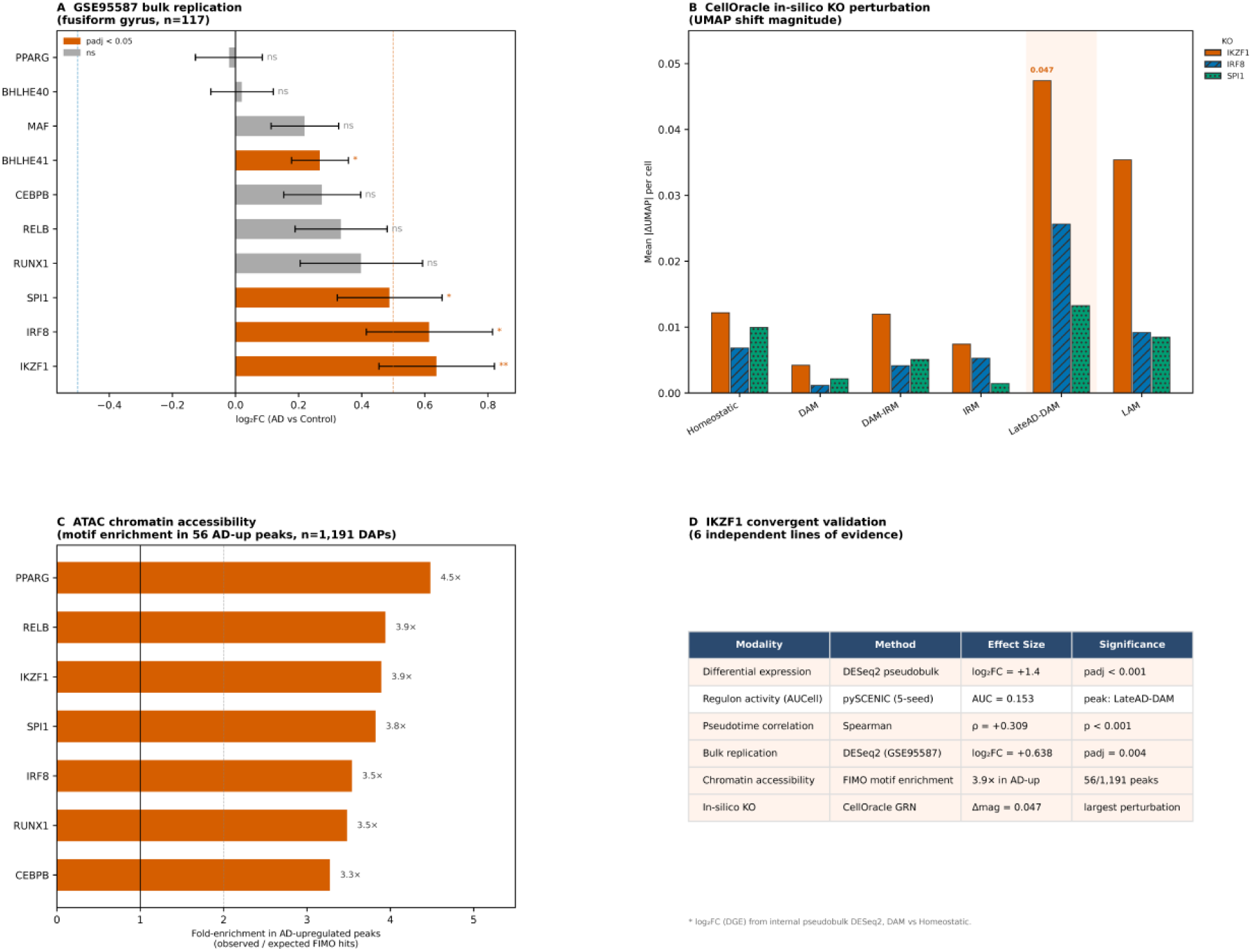
Multi-evidence prioritisation of IKZF1 as a late-disease microglial transcription-factor candidate. **(A)** Forest plot of log₂FC (AD vs control) for target TFs in GSE95587 (n = 117); *IKZF1* ranks highest among significant TFs (log₂FC = +0.638, adjusted P value =.004). **(B)** CellOracle in-silico KO: mean UMAP displacement per substate for *IKZF1*, *IRF8*, and *SPI1* KO; *IKZF1* KO produces the largest LateAD-DAM displacement (|ΔUMAP| = 0.047). **(C)** ATAC motif enrichment: fold enrichment of TF motif hits in 56 AD-upregulated open chromatin peaks; IKZF1 enriched 3.9×. **(D)** *IKZF1* multi-evidence summary table: six evidence dimensions with modality, source, method, effect size, and significance.

### Cell–cell communication identifies a neuron→microglia signalling axis not prominently described in prior AD literature

CellChat analysis (45,000 MTG nuclei; 15 cell types; 1,280 L-R pairs) identified 3,718 supported interactions; 129 involved Microglia as receiver. The top pathway, SLIT2→ROBO2 (sum_prob = 0.318), was sent by multiple neuron subtypes (Pvalb, Lamp5 Lhx6, Chandelier, Sst, Lamp5, and ExcNeuron) and has not been prominently described in prior AD microglia literature; it is an MTG-specific computational prediction requiring cross-dataset and experimental validation before any novelty claim can be made [31] (**Figure 4**; **Supplementary Fig. S6;** see **Supplementary Methods SM4** for PDF rendering notes on the chord diagram). These preliminary L-R networks often lack protein-level corroboration and are considered hypothesis-generating until validated by biophysical or functional assays [32,33]. GAS6-MERTK (sum_prob = 0.229) linked Chandelier and Pvalb interneurons to microglial efferocytosis, consistent with MERTK regulation by IRF8 [34]. Additional top pathways included SPP1 autocrine/paracrine integrin loop (sum_prob = 0.176; Microglia and Oligodendrocyte→Microglia), TGFb astrocyte/autocrine→microglia brake (0.169) —a critical checkpoint for maintaining homeostatic quiescence [35] and promoting neuronal stability [36], and COMPLEMENT autocrine (0.082).

**Figure 4.**
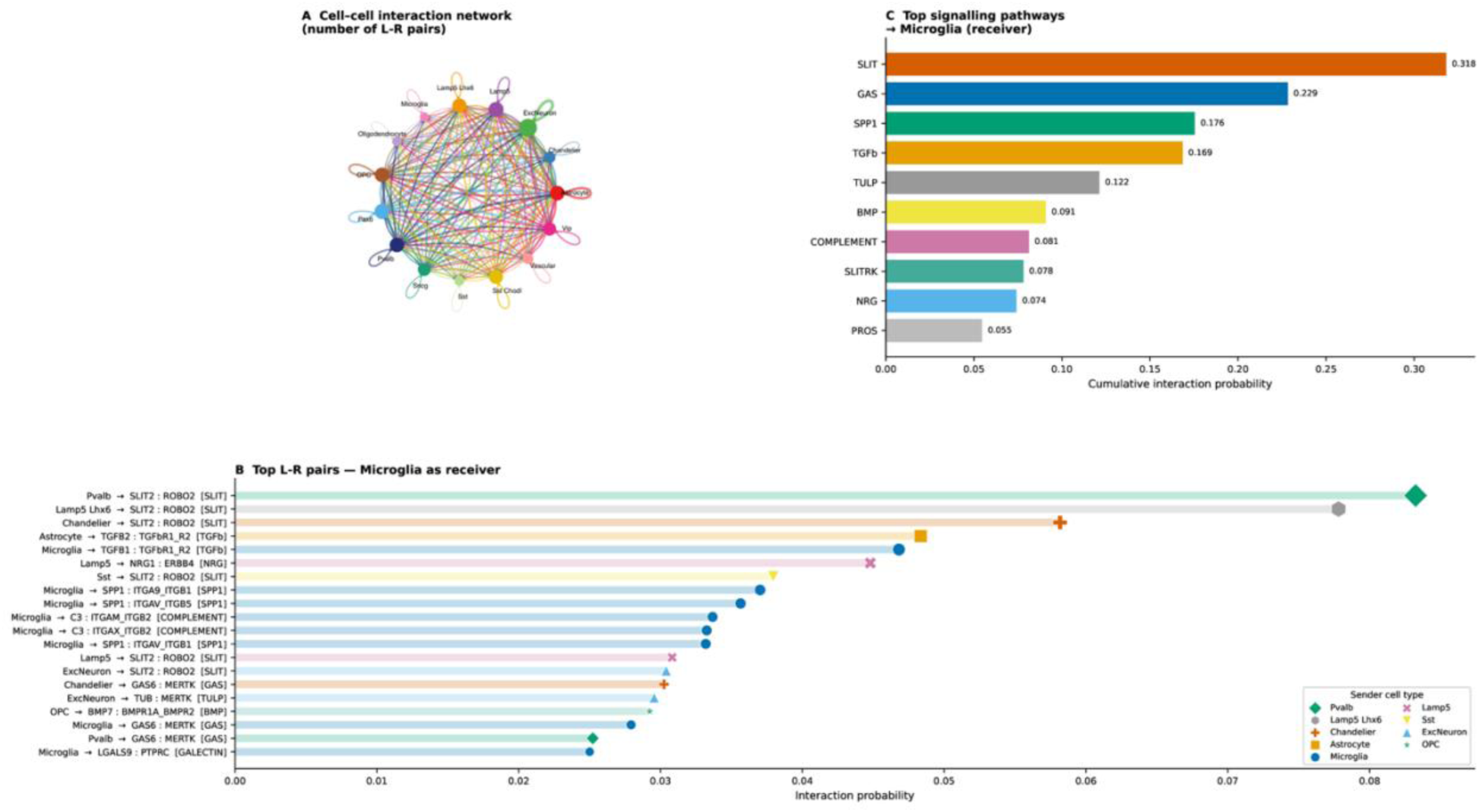
CellChat analysis identifies *SLIT2*→*ROBO2* as the top-ranked predicted interneuron–microglia signalling pathway in the SEA-AD MTG. (A) Chord diagram of cell–cell interactions across 15 MTG cell types; Microglia highlighted as receiver. **(B)** Top 20 L-R pairs with Microglia as receiver, ranked by interaction probability; sender cell type colour-coded and shape-coded. **(C)** Top 10 signalling pathways to Microglia ranked by cumulative interaction probability; *SLIT* ranks first (0.318).

### Structure-based virtual screening generates low-confidence computational hypotheses for tafamidis and diflunisal as repurposing candidates

This section is included as a methodological demonstration only; no compound meets criteria for translational advancement without experimental confirmation. Druggability assessment (fpocket) ranked PPARG (drug_score = 0.996), IRF8 AF2 DBD (0.659), and MAF bZIP (0.432) as druggable; IKZF1 ZF1–4 domain (drug_score = 0.001; ZF2 degron targeted by thalidomide analogs) was redirected to the CRBN molecular glue track (**Supplementary Fig. S7**), utilizing the established mechanism where thalidomide analogs reprogram the CRL4-CRBN E3 ligase to recruit neo-substrates like IKZF1 [37,38]. This interaction is mediated by the binding of specific C2H2 zinc finger degrons to the drug-CRBN interface [39]. Of 1,962 screened compounds (285 CNS-approved Tier-1; 1,677 Ro5/CNS-MPO-filtered Tier-2), 26 passed the dual filter (**Table 3**; **Figure 5A**). Tafamidis (CHEMBL2103837) → IRF8 was the top MM-GBSA hit (ΔG = −9.5 kcal/mol; Vina = −6.71; CNN = 0.741) but showed ligand egress in 100 ns MD (RMSD = 25.17 ± 3.56 Å last 20 ns); diflunisal (CHEMBL898) → PPARG (ΔG = −2.8 kcal/mol) showed complete egress (RMSD = 83.42 ± 2.89 Å, **Supplementary Fig. S8**). Both are transthyretin stabilisers—a post-hoc coincidence from n=2 of 1,962; no pharmacophore inference is warranted. BHLHE41 (drug_score = 0.003) and RUNX1 (all ΔG ≈ 0) are deprioritised. All six Tier-2 CRBN hits evaluated against off-target receptors (DRD2, HTR2A, hERG) failed the pre-specified selectivity threshold (SI > 2.0; observed SI < 1.0 for all six); selectivity profiling of the remaining shortlisted compounds was not performed. SPR/TSA and cell-based selectivity profiling are required before any translational inference.

**Figure 5.**
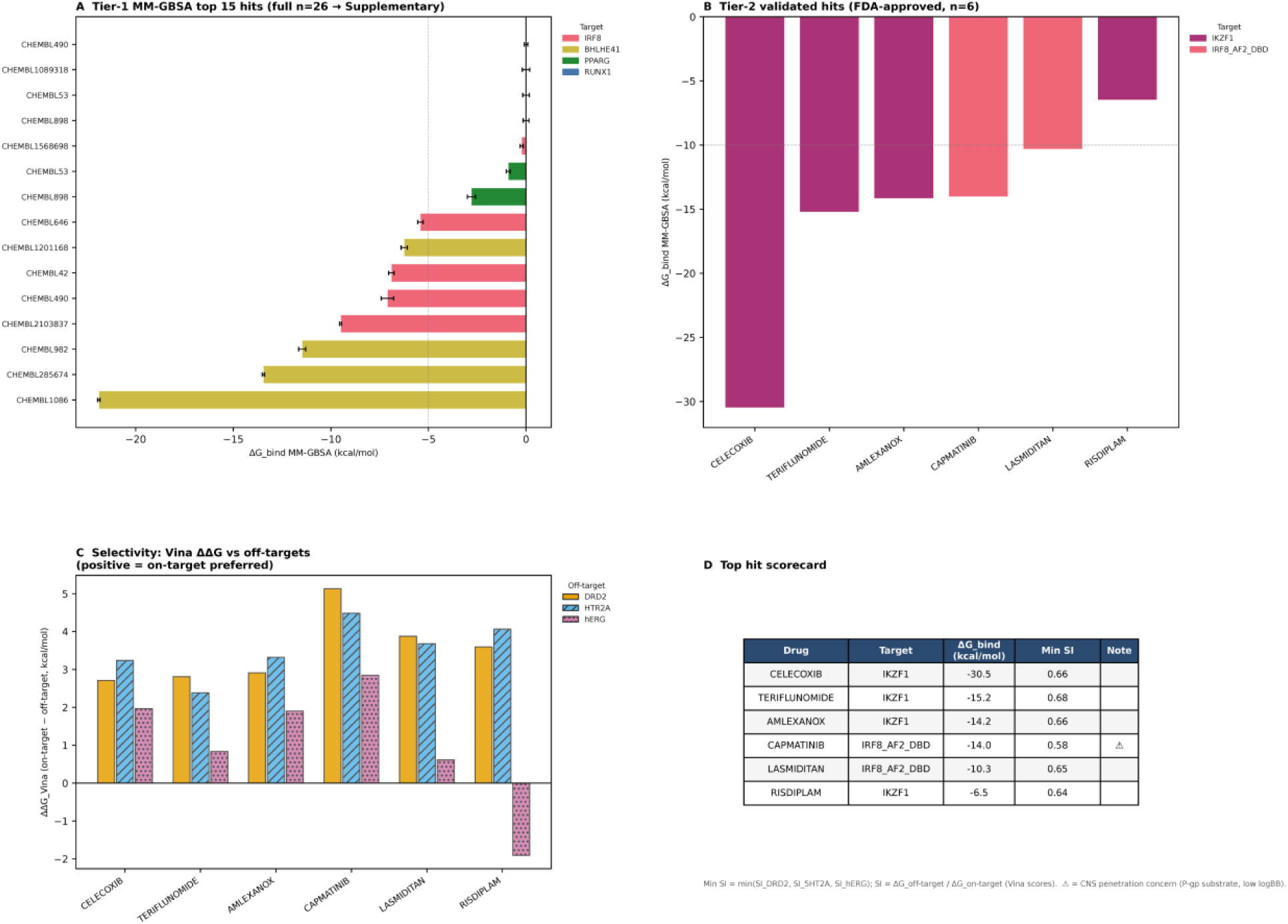
Structure-based virtual screening: consensus scoring and selectivity. Two screening tracks: Track 1 (Panel A) — orthosteric screen of 1,962 compounds against five targets (PPARG, IRF8, MAF, RUNX1, BHLHE41); Track 2 (Panels B–D) — CRBN molecular glue screen against IKZF1/IRF8 (PDB 8RQC). All compounds failed the pre-specified selectivity threshold (SI > 2.0; observed SI < 1.0 for all compounds evaluated). **(A)** MM-GBSA ΔG_bind_ (±SEM) for the top 15 Tier-1 hits; tafamidis (→IRF8) and diflunisal (→PPARG) are top-ranked. **(B)** MM-GBSA ΔG for the six Tier-2 CRBN-track hits. **(C)** Selectivity index vs DRD2, HTR2A, and hERG for the six Tier-2 compounds. **(D)** Scorecard: drug name, target, MM-GBSA ΔG_bind_, Min SI, and notes for Tier-2 hits.

**Table 3.**
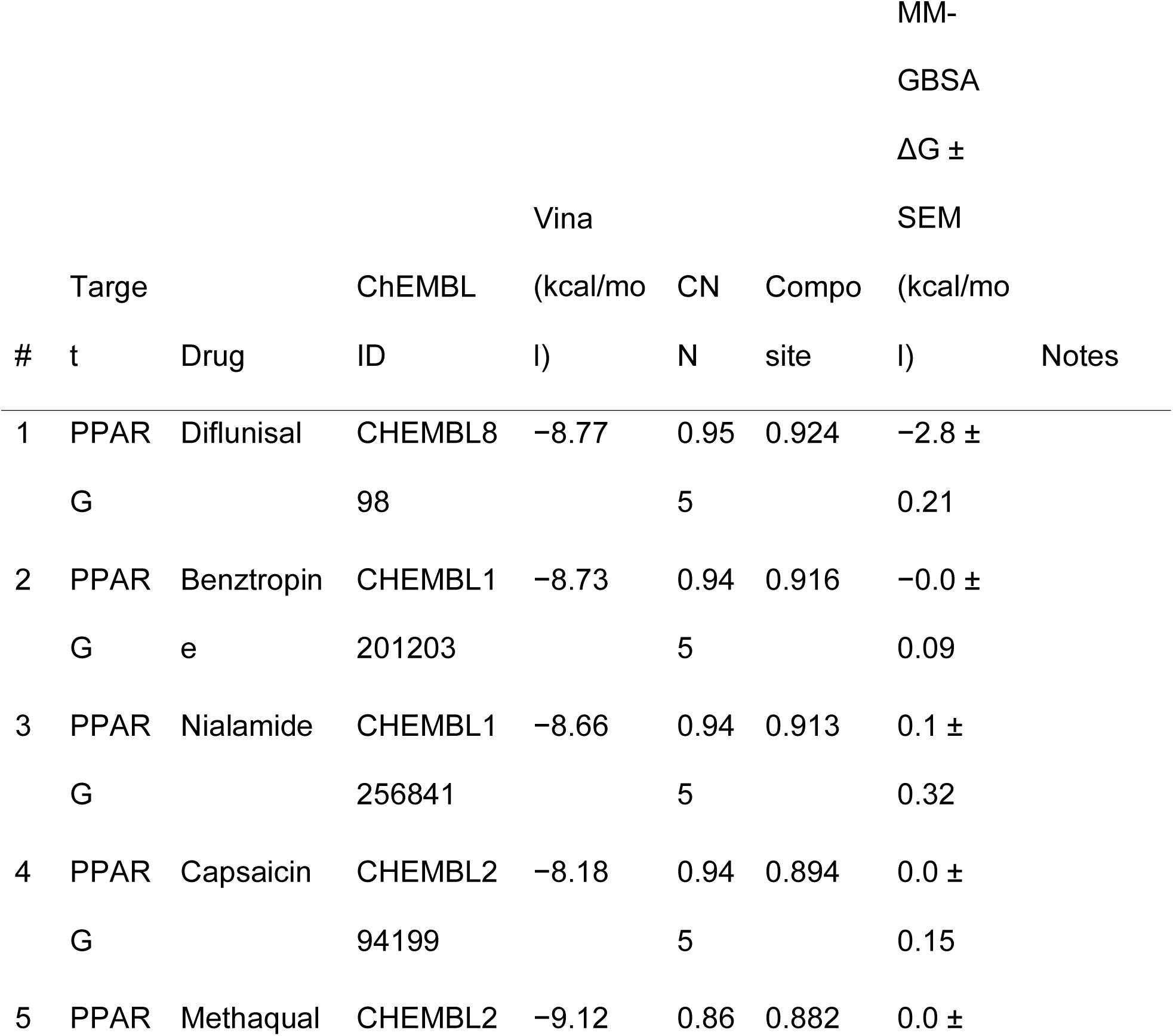
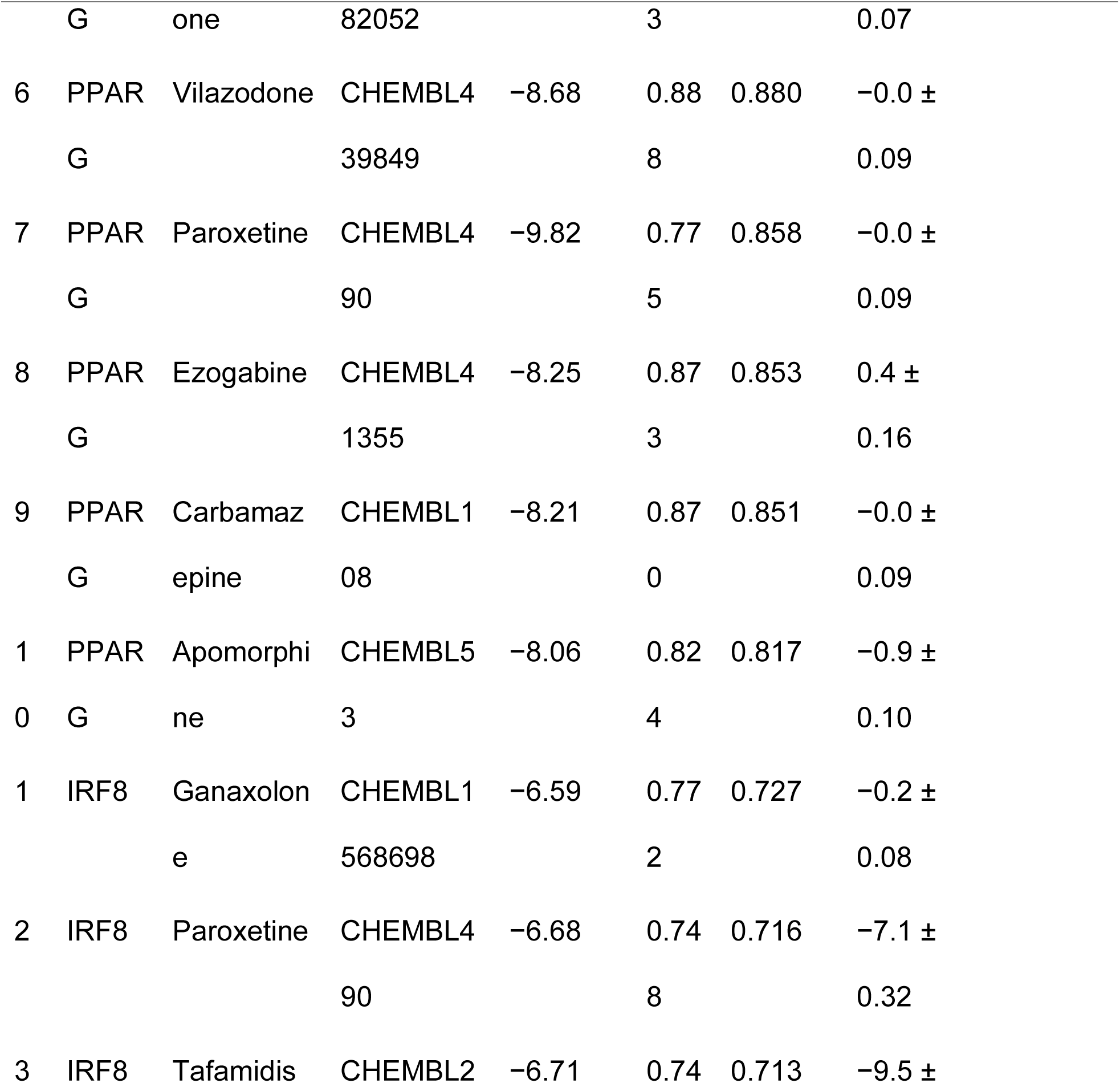
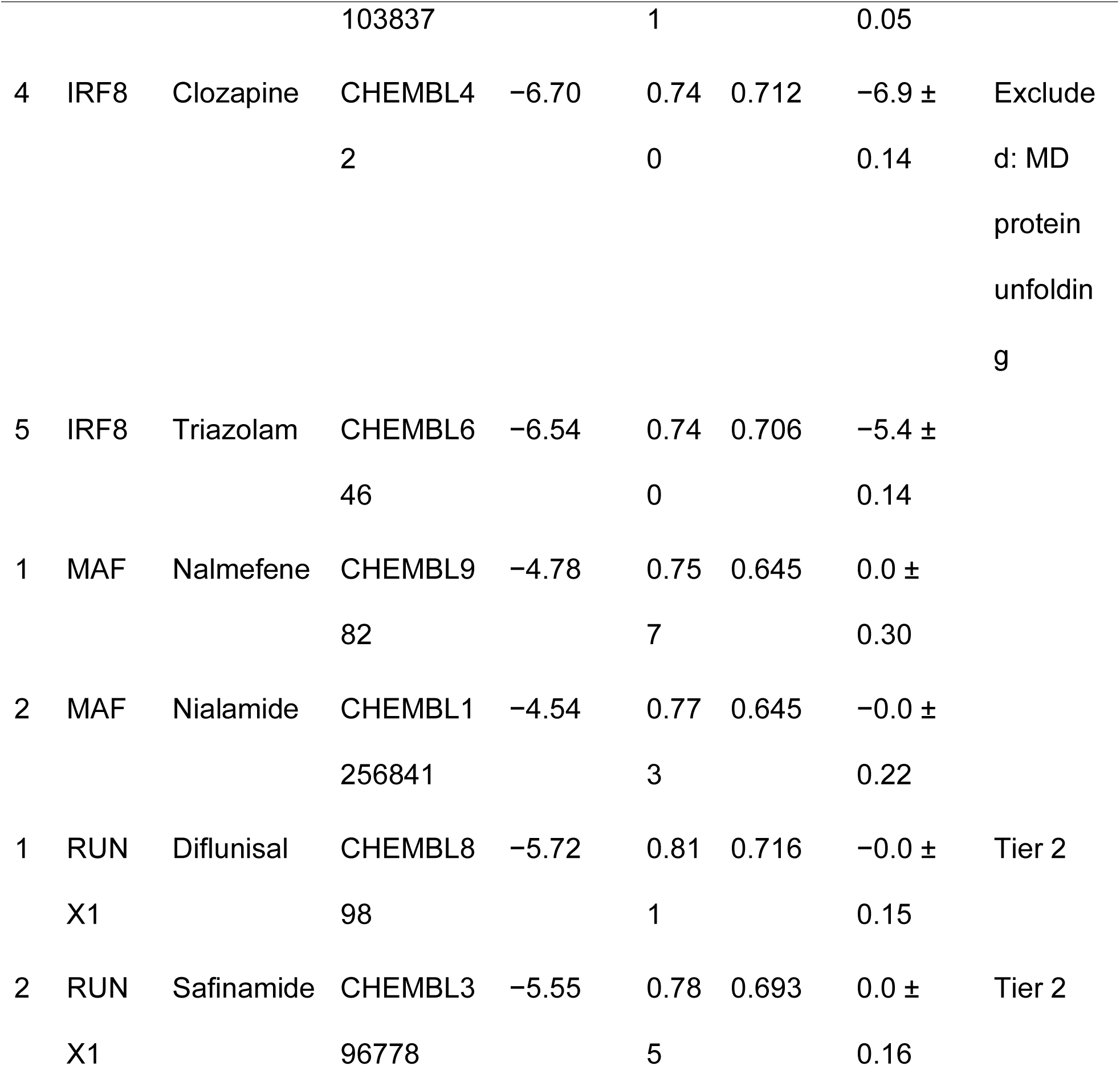
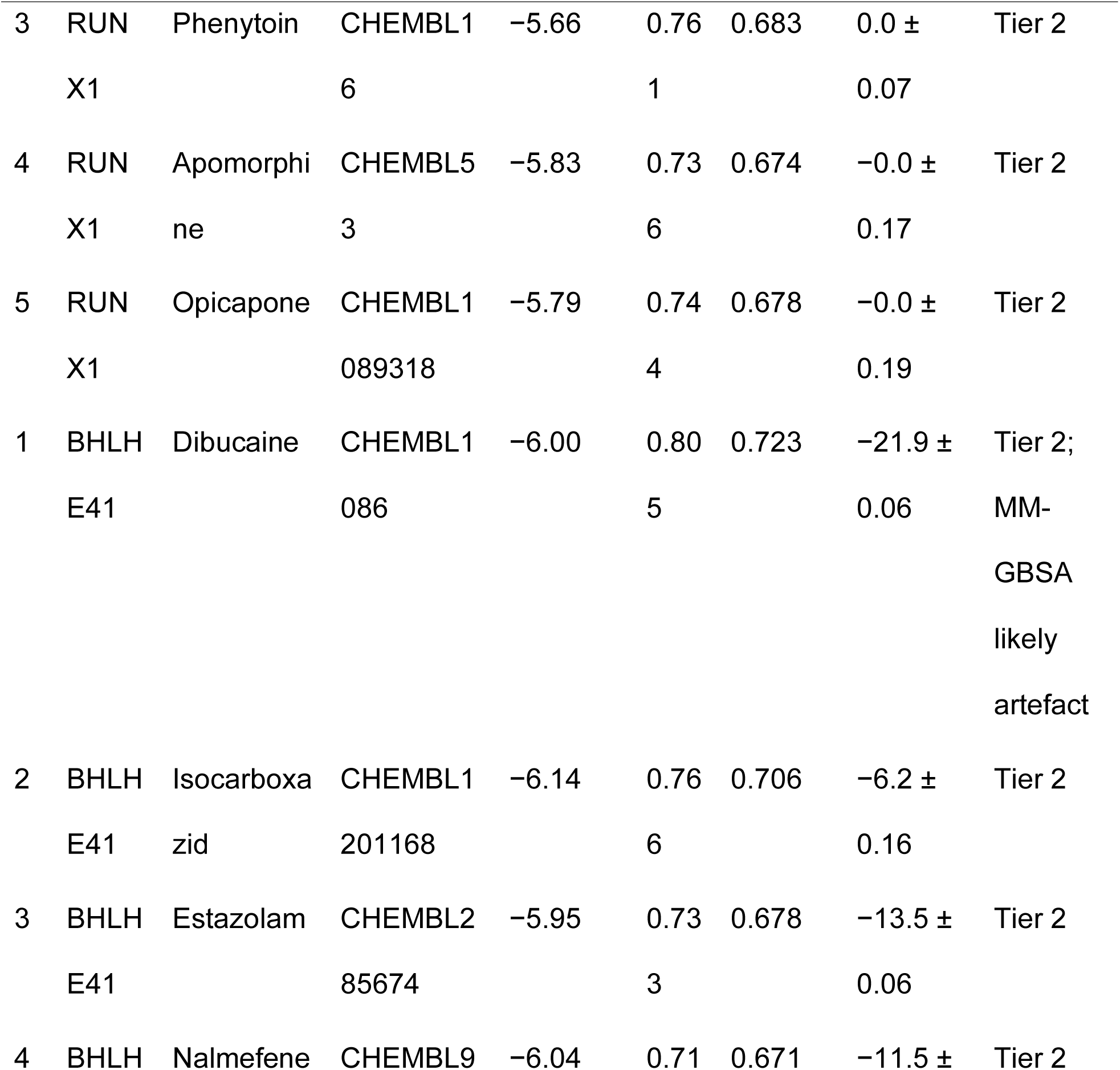
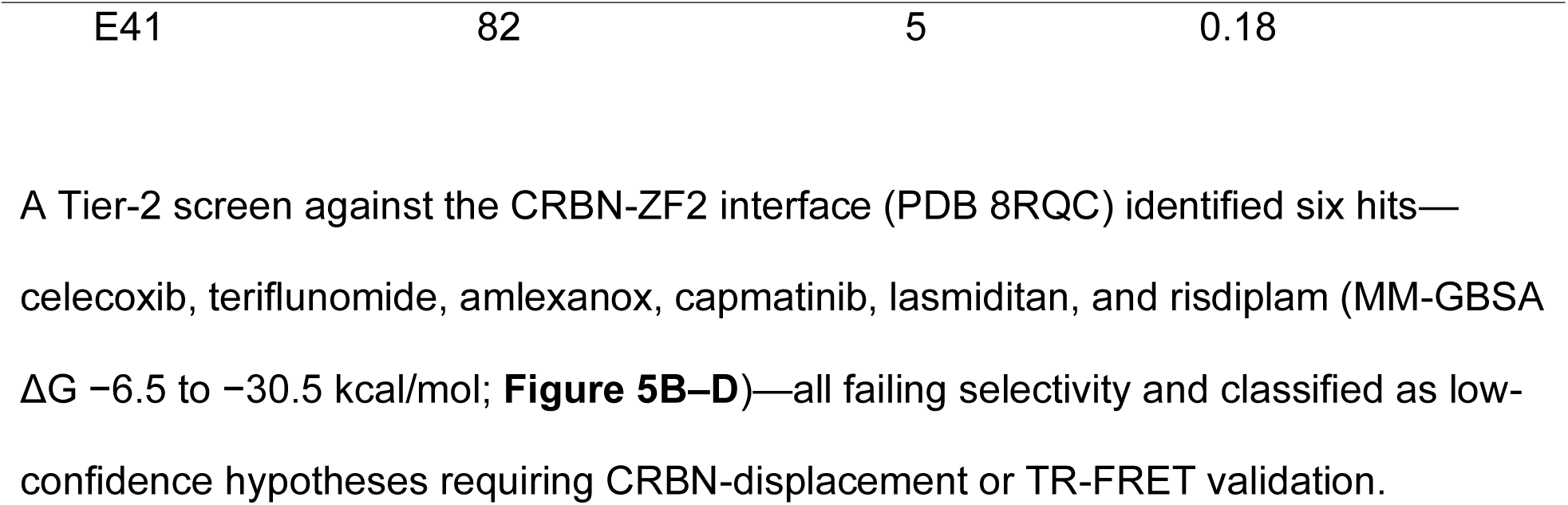
Virtual screening shortlist. . 26 compounds across five targets passing dual-filter criteria (Vina + CNN composite ≥ 0.65; Tier-2 threshold relaxed to Vina ≤ −5.5 kcal/mol for shallow-pocket targets RUNX1 and BHLHE41). MM-GBSA ΔG values near 0 kcal/mol indicate unfavourable or negligible computed binding free energy and are reported as-is. No selectivity data are shown here; selectivity profiling against DRD2, HTR2A, HERG, and DAT was performed for the six Tier-2 IRF8/IKZF1 compounds only (see Zenodo https://zenodo.org/records/19559648).

## Discussion

The central finding—that *IKZF1* is a computationally prioritised stage-specific candidate in the terminal LateAD-DAM microglial state—extends prior work by Ballasch et al. [30], who demonstrated microglial-specific *IKZF1* expression in mice, a DAM-like profile in *IKZF1*-deficient microglia, and IKZF1 protein elevation in human AD hippocampus. The present study adds: (1) stage-specific association within a human single-nucleus trajectory across 236,002 cells and 84 donors; (2) multi-modality convergence ranking of IKZF1 relative to eight other candidate TFs; and (3) nomination of the CRBN-ZF2 interface as a structural entry point via virtual screening. The SEA-AD atlas identification of *IKZF1* as an early GRN driver [10] is also prior; our contribution is the stage-resolved trajectory context and multi-modality framework applied here.

*IKZF1* encodes Ikaros, a zinc-finger transcription factor best characterised as a master regulator of lymphoid development in haematopoiesis [40,41], where it serves as a critical tumor suppressor in B-cell malignancies [42]. Its expression in tissue-resident macrophages and microglia is less studied, but emerging evidence positions IKZF1 as a regulator of innate immune quiescence: *IKZF1*-deficient microglia adopt a DAM-like transcriptional profile with increased *TREM2* and reduced *P2RY12* [30], the trajectory-resolved transition from Homeostatic to DAM states observed in our human single-nucleus analysis. Furthermore, the protein-level elevation of IKZF1 in late-stage 5xFAD mice and human AD hippocampi reported by Ballasch et al. provides biological validation for our finding that *IKZF1* regulon activity peaks in the terminal LateAD-DAM state [30]. The multi-modality convergence framework applied in this study — requiring support across GRN regulon activity, pseudotime trajectory, chromatin accessibility, independent bulk replication, and in-silico perturbation — compensates for the inherent noise of any single computational modality (SCENIC+ regulon precision ≈ 0.17–0.18 against ChIP-seq ground truth [43]) while maintaining explicit epistemic transparency about the non-independence of internal analyses.

Tafamidis (→IRF8) and diflunisal (→PPARG) are low-confidence computational hypotheses only. Both showed ligand egress in 100 ns MD; formal off-target selectivity profiling (SI > 2.0 pre-specified threshold) was not completed for these two compounds and remains a required experimental step; the MM-GBSA estimates should be treated as relative ranking scores rather than binding affinities. Both are also transthyretin stabilisers—a post-hoc coincidence from n=2 of 1,962 screened, not a biological signal. SPR or TSA with recombinant IRF8 DBD is the required experimental next step; no translational inference is warranted without experimental triage.

Separately, the cereblon (CRBN) molecular glue track identified in our virtual screen is noteworthy: approved CELMoDs (cereblon-E3 ligase modulators) including lenalidomide and pomalidomide already achieve IKZF1 protein degradation in haematopoietic malignancies. Whether this drug class could selectively modulate IKZF1 activity in microglia without systemic immunosuppression is an open question, particularly given the immunocompromised state already associated with advanced AD. The six CRBN-track hits identified here — celecoxib, teriflunomide, amlexanox, capmatinib, lasmiditan, and risdiplam — are structurally distinct from classical CELMoDs and require CRBN-displacement or TR-FRET assays to confirm cereblon engagement before further characterisation.

The SLIT2→ROBO2 signal from CellChat adds an intercellular dimension absent from most computational target-identification studies. It is an MTG-specific prediction that requires cross-dataset and experimental validation (e.g., co-culture or optogenetic perturbation) before any novelty claim; SLIT-ROBO signalling has been shown to regulate microglial process extension [44], making the prediction biologically plausible but unconfirmed. The SLIT-ROBO axis is best characterised in neuronal axon guidance during development; *ROBO2* expression on microglia may mediate process extension and chemotactic responses toward sites of neuronal injury. In AD cortex, the predominant SLIT2-expressing populations identified here — primarily inhibitory interneuron subtypes (Pvalb, Lamp5 Lhx6, Chandelier, Sst, Lamp5) with minor excitatory neuron contribution (ExcNeuron) — include cell types that are selectively vulnerable and progressively depleted with disease stage [10,45]. While functional evidence for SLIT-ROBO signaling in microglial process extension is currently limited to murine models [44], pathway-level analyses of human AD transcriptomes consistently nominate’axon guidance’ as a critical neuro-immune axis [46]. Furthermore, the reported downregulation of guidance cues during disease progression [47] mirrors the depletion of *SLIT2*-expressing interneurons observed here, suggesting that a loss of inhibitory guidance signaling may contribute to microglial dysfunction in the LateAD-DAM state. This loss, particularly within somatostatin-positive subpopulations, has been validated across independent cohorts using both single-nucleus transcriptomics and quantitative histology [48,49]. Declining *SLIT2* from dying interneurons could plausibly alter microglial process dynamics or phagocytic capacity, representing a mechanism by which neuronal loss propagates microglial dysfunction; this remains speculative.

### Limitations

All virtual screening hits, CellChat predictions, and CellOracle perturbation results are exploratory computational predictions. Internal analyses 1–5 for *IKZF1* are non-independent (same SEA-AD RNA data; not pre-registered); analysis 6 (GSE95587) is the sole external replication, in a single cohort and brain region. The absence of region-stratified pseudobulk modelling means some transcriptional differences may reflect regional composition rather than disease-state transitions. GRN inference is limited by motif database coverage: *BHLHE40*/*41* could not be evaluated because HOCOMOCO v11 lacks their atypical E-box motifs, and the *IKZF1* finding is therefore conditional on the cisTarget v10/HOCOMOCO v11 combination. The *SLIT2*→*ROBO2* pathway derives from a single brain region (MTG) and has not been replicated across the other nine SEA-AD regions. Virtual screening with implicit-solvent MM-GBSA provides relative compound rankings, not binding affinities; ligand egress in MD simulations for both top compounds underscores the necessity of biophysical triage. The SEA-AD cohort includes sex and age as covariates in differential expression models, but ancestry composition was not formally characterised; generalisability of the *IKZF1* trajectory association across ethnically diverse populations remains to be established. Sex was controlled as a covariate throughout but was not analysed as a primary biological variable; sex-stratified substate composition, sex-specific regulon dynamics, and potential sex-by-disease interactions in microglial state transitions were not examined. Given emerging evidence for sex-dimorphic microglial responses in AD, this represents a limitation and a priority for future stratified analyses.

### Generalizable prioritisation framework

The DAM-DRUG pipeline is transferable to any disease-relevant cell type with atlas-level single-nucleus data and at least one independent bulk cohort. The six-step decision logic (cell-state trajectory → TF candidate set → GRN inference → multi-modality convergence scoring → structural druggability/virtual screening → cell-cell communication context) and advancement criteria (≥4/6 evidence dimensions) are documented with substitution guidance in the GitHub repository README.

Immediate experimental priorities for validating the computational findings reported here include: (a) surface plasmon resonance (SPR) or thermal shift assay (TSA) to confirm tafamidis binding to recombinant IRF8 DBD; (b) CRISPR-mediated knockout of *IKZF1* in iPSC-derived or primary human microglia followed by transcriptomic profiling to determine whether *IKZF1* loss reproduces or accelerates the LateAD-DAM signature; (c) cross-dataset replication of the SLIT2→ROBO2 pathway in an independent snRNA-seq dataset with paired interneuron and microglia populations; and (d) region-stratified pseudobulk modelling across the ten SEA-AD brain regions to determine whether the *IKZF1* trajectory association is cortex-wide or region-specific.

## Conclusion

Applying a six-step multi-modality convergence framework to 236,002 microglial nuclei from 84 postmortem AD donors, we nominate *IKZF1* as the primary computationally prioritised transcription factor in the terminal LateAD-DAM state. This nomination rests on concordant evidence across pseudobulk differential expression, GRN regulon activity, diffusion pseudotime trajectory, chromatin accessibility motif enrichment, independent bulk RNA-seq replication (GSE95587), and in-silico perturbation — with external replication in a single independent cohort representing the sole non-SEA-AD evidence line. CellChat analysis identifies SLIT2→ROBO2, predominantly from inhibitory interneuron subtypes, as the top predicted neuron-to-microglia signalling pathway in the middle temporal gyrus, a hypothesis requiring cross-dataset and experimental validation. Structure-based virtual screening nominates tafamidis (→IRF8) and diflunisal (→PPARG) as low-confidence repurposing hypotheses; ligand egress in MD simulations and incomplete selectivity profiling preclude any translational inference without biophysical triage. Together, these findings establish a transferable computational prioritisation framework for druggable TF candidates in disease-relevant human microglial states.

## Acknowledgements

Computational analyses were performed on the TRUBA ARF cluster (TÜBİTAK ULAKBİM). SEA-AD data were generated by the Allen Institute for Brain Science.

## Author Contributions

Ç.Ö.: Conceptualization, Data curation, Formal analysis, Investigation, Methodology, Project administration, Software, Validation, Visualization, Writing – original draft, Writing – review and editing.

## Statements and Declarations

*Ethical considerations*: This study is a secondary computational analysis of publicly available, de-identified postmortem brain data (SEA-AD atlas; GSE95587); no primary human samples were collected. No IRB approval was required, consistent with standard practice for analyses of pre-existing, de-identified datasets and in accordance with the Declaration of Helsinki.

*Consent to participate*: Not applicable.

*Consent for publication*: Not applicable.

*Declaration of conflicting interest*: The author declared no potential conflicts of interest with respect to the research, authorship, and/or publication of this article.

*Funding statement*: This research did not receive any specific grant from funding agencies in the public, commercial, or not-for-profit sectors.

*Data availability*: All analysis scripts are available at https://github.com/cagriozkurt/DAM-DRUG. Supplementary data files are deposited at Zenodo (https://zenodo.org/records/19559648). The SEA-AD microglial dataset is freely available from the AWS open data registry (s3://sea-ad-single-cell-profiling; https://registry.opendata.aws/allen-sea-ad-atlas). GSE95587 is available from NCBI GEO. Compound structures were obtained from ChEMBL (max_phase = 4). AlphaFold2 structures were obtained from the EBI AlphaFold Database (v6). Experimental PDB structures are available from the RCSB Protein Data Bank.

