## Supplementary Methods and Tables S1-S7 for "Single-cell gene networks nominate *IKZF1* as an Alzheimer’s microglial regulator"

Çağrı Özkurt[1]

[1] Department of Pharmacology, Faculty of Pharmacy, Hacettepe University, 06230 Ankara, Turkey

### Table of Contents

1. Supplementary Methods
2. Supplementary Tables
   - Table S1. SEA-AD Supertype-to-Functional-Substate Mapping
   - Table S2. Analytical Pipeline and Donor Inclusion Criteria
   - Table S3. Virtual Screening Parameters and Targeted Structural Domains
   - Table S4. Pre-specified Thresholds and Consensus Scoring Metrics
   - Table S5. Inventory of Stochastic Seeds and Algorithmic Parameters for Pipeline Reproducibility
   - Table S6. Microglial Substate Cell Counts by Brain Region
   - Table S7. GRNBoost2 Random Seeds and Edge Counts
3. Supplementary Figures
   - Figure S1. Quality control metrics and dataset overview
   - Figure S2. BHLHE40/41 motif-agnostic pseudobulk co-expression analysis
   - Figure S3. SLIT2 and ROBO2 expression across MTG cell types
   - Figure S4. pySCENIC regulon activity heatmap
   - Figure S5. CellOracle in-silico TF knockout perturbation magnitudes
   - Figure S6. Microglial substate UMAPs (faceted)
   - Figure S7. AlphaFold2 per-residue pLDDT quality scores
   - Figure S8. Ligand RMSD traces from 100 ns MD simulations

### 1. Supplementary Methods

#### *SM1. GRNBoost2 Random Seeds*

GRNBoost2 is a stochastic gradient-boosted regression tree algorithm for co-expression network inference whose output varies with the random seed. To reduce seed-dependent noise, five independent GRN runs were performed; edges appearing in ≥ 3/5 seeds were retained for downstream RcisTarget pruning. **Supplementary Table S7** lists the five seeds, the number of edges per run, and the final consensus edge count.

The seed=42 run was executed via code/slurm/06_run_pySCENIC.slurm (initial run); seeds 1–4 were added via code/slurm/07_grn_multiseed.slurm (SLURM array job). Edge aggregation was performed by code/phase2_GRN/02_aggregate_grn.py. No systematic differences in regulon composition were detected across seeds (all five seed runs recovered the same 46 cisTarget-pruned regulons under the ≥ 3/5 consensus rule), confirming that the IKZF1 regulon identification is not an artefact of a single seed.

#### *SM2. Motif Database Choice (HOCOMOCO v11 / cisTarget v10)*

The cisTarget v10 rankings database and motif annotation table (motifs-v10nr_clust-nr.hgnc-m0.001-o0.0.tbl) were selected because they represent the most comprehensive non-redundant human TF motif collection available within the pySCENIC framework at the time of analysis (2024–2025), and because they include HOCOMOCO v11 entries with HGNC gene annotation, which is required for mapping motifs to the human TF target list used in this study.

HOCOMOCO v11 justification. HOCOMOCO v11 provides high-quality position weight matrices derived from ChIP-seq experiments for 769 human TFs. The IKZF1 zinc-finger binding motif is represented in HOCOMOCO v11 (model IKZF1_HUMAN.H11MO.0.B), enabling RcisTarget to recover the IKZF1(+) regulon. By contrast, BHLHE40 and BHLHE41 use atypical E-box variants not captured by existing HOCOMOCO v11 matrices (the canonical *BHLHE40*/*41* E-box motif CACGCG is represented only at low specificity), explaining their absence from the 46 cisTarget-pruned regulons. This is a database coverage limitation, not a GRN inference failure.

Alternative databases. Repeating the RcisTarget pruning step with JASPAR 2024 (which contains an updated *BHLHE40* matrix based on expanded ChIP-seq data) may partially recover *BHLHE40*/*41* regulons and is proposed as a future validation. The current results should be interpreted as specific to the cisTarget v10 / HOCOMOCO v11 combination.

#### *SM3. scMultiomeGRN Pivot to ATAC Differential Accessibility Validation*

The original analysis plan included a scMultiomeGRN analysis (paired RNA+ATAC single-cell regulatory network inference). This was pivoted during analysis after discovering that the SEA-AD MTG RNA object (SEAAD_MTG_RNAseq_final-nuclei.2024-02-13.h5ad) and the MTG ATAC object (SEAAD_MTG_ATACseq_final-nuclei.2024-12-06.h5ad) share only approximately 1.3% of cell barcodes — a fundamental incompatibility with the paired-multiome requirement of scMultiomeGRN, which requires the same cells to be profiled by both modalities.

The 1.3% overlap arises because the SEA-AD atlas was generated by the 10x Multiome protocol (RNA + ATAC from the same nuclei), but the two modalities were separately preprocessed and released as independent AnnData objects with different cell filtering criteria; barcodes retained after RNA QC do not fully overlap with those retained after ATAC QC. For scMultiomeGRN, the expected overlap would be ≥ 90% for a properly paired multiome release; 1.3% is consistent with independent QC filtering and is not a data download error.

Consequently, the ATAC data were repurposed as an orthogonal chromatin accessibility validation rather than a paired GRN input: differential accessibility testing (Severely Affected vs non-Severely-Affected donors; 218,882 peaks tested; Mann-Whitney U, BH FDR) was used to identify AD-upregulated peaks, which were then scanned for TF motif enrichment by FIMO (see Section 2.5 in Methods). This analysis provides corroborative rather than independent evidence for IKZF1 (same donors, different assay).

#### *SM4. Reproducibility Notes and Known Rendering Issues*

**Figure 4A** (chord diagram) — blank rendering in some PDF viewers. The chord diagram in Panel A of the CellChat figure (results/figures/fig_cellchat.pdf) may appear blank in PDF viewers that do not support the transparency and gradient fills generated by the R circlize package. This is a rendering limitation of certain PDF engines, not an analysis error. The PNG version (results/figures/fig_cellchat.png) renders correctly in all tested environments. Additionally, converting the PDF to PNG using pdftoppm requires the Poppler library (poppler-utils on Ubuntu/Debian); if pdftoppm is unavailable, the PDF can be opened directly in a browser (e.g., Firefox, Chrome) or viewed using Evince or Okular on Linux.

GRNBoost2 non-determinism between platforms. GRNBoost2 (via pySCENIC grn) uses DAAL (Intel oneDAL) for tree operations on x86_64. The same random seed may produce slightly different edge weights on ARM vs x86_64 hardware due to floating-point instruction ordering differences. The TRUBA cluster uses x86_64 (Intel/AMD). Results may not be exactly reproduced on ARM hardware (e.g., Apple M-series) even with identical seeds; however, the 5-seed consensus criterion substantially reduces seed-level variance.

Container images and exact software versions. The following Apptainer/Singularity images were used for all analyses. Image digests as pulled from GHCR are not pinned in the SLURM scripts (the :latest tag is used); exact image digests at pull time are not recorded. Future reproducibility requires pinning to a specific image digest or rebuilding from the Dockerfiles provided in docker/:

| Container SIF | Docker image | Key environments |
|---|---|---|
| containers/scenic.sif | ghcr.io/cagriozkurt/dam-drug-scanpy:latest | scanpy_env (Python, Scanpy, pyDESeq2, |

| Container SIF | Docker image | Key environments |
|---|---|---|
| | | CellOracle), scenic (pySCENIC 0.12.1) |
| containers/fpocket-env.sif | ghcr.io/cagriozkurt/dam-drug-fpocket:latest | scanpy_env + fpocket 4.x, PDBFixer, OpenMM |
| containers/cellchat.sif | ghcr.io/cagriozkurt/dam-drug-cellchat:latest | R 4.3, CellChat v2 |

### 2. Supplementary Tables

***Supplementary Table S1. SEA-AD Supertype-to-Functional-Substate Mapping***

Pre-annotated Supertype labels from the SEA-AD multi-regional microglial release were mapped to six functional substates by direct lookup. The "-SEAAD" suffix reflects the annotation tier present in the SEA-AD multi-regional h5ad release. Non-suffixed variants (e.g. Micro-PVM_2_3) were absent from this release and are not included in the mapping.

| SEA-AD Supertype label | Functional substate | Abbreviation | n cells |
|---|---|---|---|
| Micro-PVM_1 | Homeostatic | HM | 7,909 |
| Micro-PVM_2 | Disease-associated microglia | DAM | 141,248 |
| Micro-PVM_2_3-SEAAD | DAM-IRM (transition state) | DAM-IRM | 53,815 |
| Micro-PVM_3-SEAAD | Interferon-responsive microglia | IRM | 29,805 |
| Micro-PVM_2_1-SEAAD | Late-AD DAM | LateAD-DAM | 1,458 |
| Micro-PVM_4-SEAAD | Lipid-associated microglia | LAM | 1,767 |

Mapping was implemented in code/phase5_validation/01_celloracle_perturbation.py and applied consistently across all pipeline modules. DAM-IRM was retained as a distinct substate in trajectory, AUCell, and CellOracle analyses but excluded from primary DGE contrasts owing to its hybrid co-expression profile (TREM2/APOE + IFIT1/MX1).

***Supplementary Table S2. Analytical Pipeline and Donor Inclusion Criteria for the Multi-Modality Framework.*** *This table details the specific data objects, donor counts, and filtering logic applied across each computational module of the study. Of the 84 total donors in the SEA-AD multi-regional atlas, all individuals contributed to the core trajectory and cell-cell communication analyses. However, stricter inclusion thresholds were applied to downstream steps to ensure statistical robustness: pseudobulk differential gene expression (DGE) required a minimum of 10 cells per donor-state, resulting in the exclusion of the sparse LateAD-DAM state from donor-level testing. The ATAC-seq module utilized a subset of 84 donors stratified by severe neuropathology for chromatin accessibility mapping.*

| **Analysis** | **Dataset object** | **Donors included** | **Exclusion criterion** |
|---|---|---|---|
| Trajectory inference (PAGA + DPT) | Multi-regional RNA | All 84 | None |
| Pseudobulk DGE (DESeq2) — DAM, IRM | Multi-regional RNA | 84 / 84 | ≥10 cells/donor in case state |
| Pseudobulk DGE (DESeq2) — LAM | Multi-regional RNA | 27 / 84 | ≥10 cells/donor; 57 donors have <10 LAM cells |
| Pseudobulk DGE (DESeq2) — LateAD-DAM | Multi-regional RNA | 0 / 84 | Excluded entirely; median 4 cells/donor |
| pySCENIC GRN inference (100K subset) | Multi-regional RNA | All 84 (proportional) | None; simple random draw without replacement; no donor-balance correction |
| AUCell scoring + pseudotime correlation | Multi-regional RNA | All 84 (proportional) | Same 100K subset as GRNBoost2 |
| ATAC differential accessibility | MTG ATAC | 84 of 86 (2 excluded: no Severely Affected label) | Severely Affected (n=11) vs non-Affected (n=73); no per-donor cell minimum |
| CellChat L-R analysis | MTG RNA | All 84 with MTG data | None beyond min.cells=10 per cell type per interaction |
| CellOracle in-silico | Multi-regional RNA | All 84 (proportional) | 20K stratified subsample |

| Analysis | Dataset object | Donors included | Exclusion criterion |
|---|---|---|---|
| KO | | | (proportional allocation, floor 200 cells/state);; no donor exclusion |
| BHLHE40/41 co-expression rescue | Multi-regional RNA | 84 pseudobulk samples | Same as pseudobulk DGE |
| GSE95587 bulk replication | External (GEO) | 84 AD / 33 control (n=117) | Pre-existing study design; no additional exclusions by us |

***Supplementary Table S3. Virtual Screening Parameters and Targeted Structural Domains.*** *The following table details the grid box coordinates, pocket dimensions, and druggability metrics utilized for the structure-based virtual screening of 1,962 compounds against six prioritized microglial transcription factor (TF) domains.*

| Target | PDB | Pocket definition | Center x, y, z (Å) | Box (Å) | fpocket vol (Å³) | drug_score |
|---|---|---|---|---|---|---|
| PPARG | 1FM9 | fpocket pocket1 centroid | 12.487, −3.853, 46.330 | 24 | 1324.5 | 0.996 |
| IRF8 | AF2 DBD | fpocket pocket1 centroid | −29.595, −2.566, 20.373 | 20 | 499.7 | 0.659 |
| MAF | 4EOT | fpocket pocket1 centroid | −43.648, 14.275, −20.945 | 18 | 231.6 | 0.432 |
| RUNX1 | 1LJM | fpocket pocket1 centroid | −7.658, 38.668, 12.552 | 20 | 486.6 | 0.092 |
| BHLHE41 | AF2 bHLH | fpocket pocket1 centroid | −48.990, 13.505, 34.681 | 20 | 402.0 | 0.003 |
| IKZF1 | 8RQC | QFC ligand centroid | 0.566, −2.235, 4.760 | 20 | — | — |

***Supplementary Table S4. Pre-specified Thresholds and Consensus Scoring Metrics for Virtual Screening.*** *The table defines the multi-layer decision logic used to shortlist candidate compounds for each transcription factor (TF) target based on structural druggability and docking performance. Vina thresholds were adjusted according to pocket topology, with more stringent criteria (≤ −6.5 kcal/mol) applied to deep, well-defined pockets (PPARG, IRF8) and relaxed criteria (≤ −5.5 kcal/mol) for shallow protein-protein interface surfaces (RUNX1, BHLHE41). The CNN threshold (≥0.70) was held constant across targets to ensure high-confidence pose quality. Note that IKZF1 was excluded from orthosteric screening due to the absence of a druggable pocket and redirected to a molecular glue/PROTAC prioritization track.*

| Target | drug_score | Vina threshold (kcal/mol) | CNN threshold | Rationale |
|---|---|---|---|---|
| PPARG | 0.996 | ≤ −6.5 | ≥ 0.70 | Deep, well-defined LBD; −6.5 kcal/mol is a standard Vina “hit” cutoff approximating low-μM affinity |
| IRF8 | 0.659 | ≤ −6.5 | ≥ 0.70 | Defined hydrophobic DBD groove; same stringent criterion as PPARG |
| RUNX1 | 0.092 | ≤ −5.5 | ≥ 0.70 | Shallow Runt-domain surface; Vina systematically underscores shallow interfaces; relaxed to −5.5 to retain candidates for CNN rescoring |
| BHLHE41 | 0.003 | ≤ −5.5 | ≥ 0.70 | Same reasoning as RUNX1; bHLH surface not deeply druggable |
| MAF | 0.432 | none | ≥ 0.75 | Best GNINA-rescored Vina score was |

| Target | drug_score | Vina threshold (kcal/mol) | CNN threshold | Rationale |
|---|---|---|---|---|
| | | | | −5.29 kcal/mol (top-30 Vina compounds rescored);no compound meets any realistic Vina threshold after CNN rescoring; CNN rescoring used as sole criterion with slightly raised cutoff (0.75) |
| IKZF1 | — | n/a | n/a | No druggable orthosteric pocket; CNS library pharmacophores incompatible with CRBN-interface molecular glue chemistry; excluded from orthosteric screen |

***Supplementary Table S5. Inventory of Stochastic Seeds and Algorithmic Parameters for Pipeline Reproducibility.*** *The table lists the fixed random seeds and computational parameters applied across the multi-stage DAM-DRUG workflow to ensure the reproducibility of stochastic analyses. Core trajectory inference (KNN, and PAGA) utilized a global seed (s=42) to stabilize the latent space and pseudotime ordering of the 236,002 microglial nuclei. The scVI latent space was loaded from the pre-processed SEA-AD atlas (scVI run by the SEA-AD consortium; seed not set by this pipeline). For Gene Regulatory Network (GRN) inference, a multi-seed consensus strategy was employed (seeds 42, 1, 2, 3, and 4) to address the inherent stochasticity of GRNBoost2; edges were retained only if present in ≥3 of the 5 independent runs. Downstream simulations, including CellOracle in silico knockouts and CellChat intercellular communication, were fixed with seed 42 to ensure consistent cell subsampling and transition probability estimates. Molecular Dynamics (MD) simulations* *used system-generated random seeds (gen_seed=−1) for velocity generation, with the resulting 100 ns trajectories archived for audit.*

| Analysis | Function / parameter | Seed value(s) |
|---|---|---|
| KNN graph (trajectory) | sc.pp.neighbors(random_state=) | 42 |
| PAGA-initialised UMAP | sc.tl.umap(init_pos="paga", random_state=) | 42 |
| pySCENIC 100K subsample | sc.pp.subsample(random_state=) | 42 |
| pySCENIC GRNBoost2 | pyscenic grn --seed | 42, 1, 2, 3, 4 (5 independent runs) |
| ATAC background peak sampling | DataFrame.sample(random_state=) | 42 |
| CellChat MTG subsampling (5K/type) | np.random.seed() | 42 |
| CellOracle stratified subsample | np.random.default_rng() | 42 |
| GROMACS MD velocity generation | gen_seed | −1 (system random; trajectories archived on TRUBA) |
| GNINA re-docking | --seed | 42 |

***Supplementary Table S6. Microglial Substate Cell Counts by Brain Region***

| Brain region | Homeostatic | DAM | DAM-IRM | IRM | LateAD-DAM | LAM | Total |
|---|---|---|---|---|---|---|---|
| AnG | 454 | 5,858 | 2,050 | 1,422 | 53 | 271 | 10,108 |
| DFC | 1,190 | 25,056 | 9,173 | 4,817 | 360 | 203 | 40,799 |
| FI | 860 | 12,337 | 2,518 | 1,289 | 60 | 42 | 17,106 |
| HIP | 508 | 8,369 | 2,147 | 2,793 | 36 | 79 | 13,932 |
| ITG | 516 | 8,631 | 4,444 | 1,945 | 171 | 114 | 15,821 |
| LEC | 283 | 8,485 | 1,490 | 1,295 | 57 | 40 | 11,650 |
| MEC | 1,662 | 36,241 | 15,573 | 7,721 | 238 | 281 | 61,716 |
| MTG | 1,383 | 21,090 | 8,278 | 4,468 | 282 | 406 | 35,907 |
| STG | 515 | 7,863 | 4,434 | 1,942 | 150 | 85 | 14,989 |
| V1C | 538 | 7,318 | 3,708 | 2,113 | 51 | 246 | 13,974 |
| Total | 7,909 | 141,248 | 53,815 | 29,805 | 1,458 | 1,767 | 236,002 |

AnG = angular gyrus; DFC = dorsolateral prefrontal cortex; FI = frontal insula; HIP = hippocampus; ITG = inferior temporal gyrus; LEC = lateral entorhinal cortex; MEC = medial entorhinal cortex; MTG = middle temporal gyrus; STG = superior temporal gyrus; V1C = primary visual cortex.

***Supplementary Table S7. GRNBoost2 Random Seeds and Edge Counts***

| Run | Seed | Edges in adjacency matrix | Edges in ≥ 3/5 consensus |
|---|---|---|---|
| 1 | 42 | — | — |
| 2 | 1 | — | — |
| 3 | 2 | — | — |
| 4 | 3 | — | — |
| 5 | 4 | — | — |
| Consensus | ≥ 3/5 seeds | — | 1,458,113 |

Note: Per-run edge counts are recorded in results/phase2/GRN/adj_matrix_seed{1..4}.tsv and results/phase2/GRN/adj_matrix.tsv; exact counts derivable by wc -l on each file. The consensus was produced by code/phase2_GRN/02_aggregate_grn.py.

#### 3. Supplementary Figures

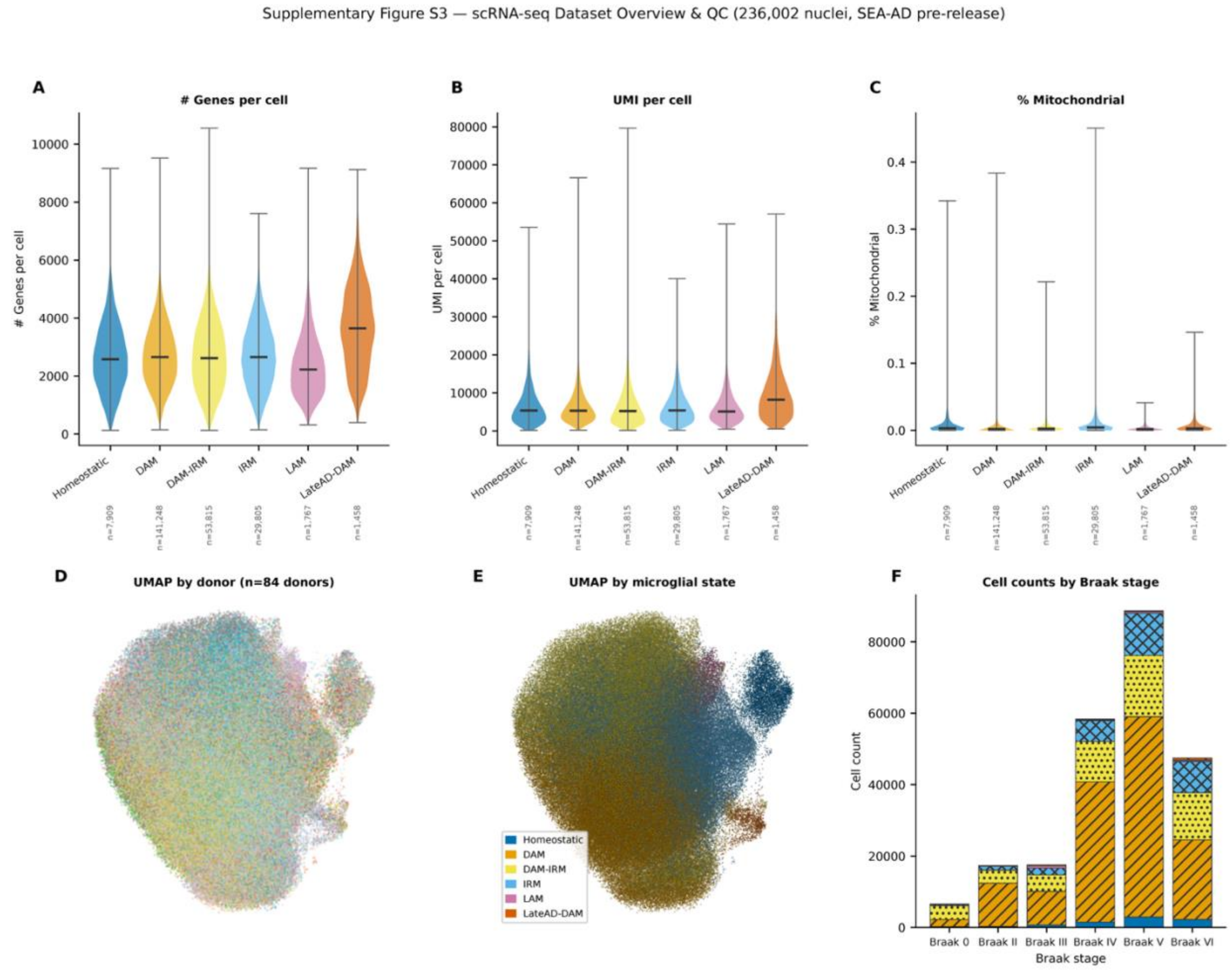


***Supplementary Figure S1. Quality control metrics and dataset overview for the SEA-AD microglial dataset (236,002 nuclei).***

(A) Violin plots of detected genes per nucleus across six microglial substates; median indicated by horizontal bar. (B) Violin plots of UMI counts per nucleus across substates. (C) Violin plots of mitochondrial gene fraction per nucleus across substates; values near zero confirm effective removal of low-quality nuclei by the Allen Institute pre-processing pipeline. (D) UMAP embedding coloured by donor identity (n = 84 donors); inter-donor mixing confirms effective batch integration by the pre-release scVI pipeline. (E) UMAP coloured by microglial substate with thin semi-transparent point borders to aid discrimination of overlapping clusters. (F) Stacked bar chart of cell counts per microglial substate across Braak stages 0, II–VI, with distinct hatch patterns per substate for accessibility; DAM and LateAD-DAM are progressively enriched at higher Braak stages.

QC was performed by the Allen Institute BICAN/SEA-AD pipeline (Scrublet doublet removal, scVI latent-space batch integration); no additional QC filtering was applied in this study. The low mitochondrial fraction across all substates (Panel C) confirms effective nucleus isolation and removal of damaged cells prior to data release.

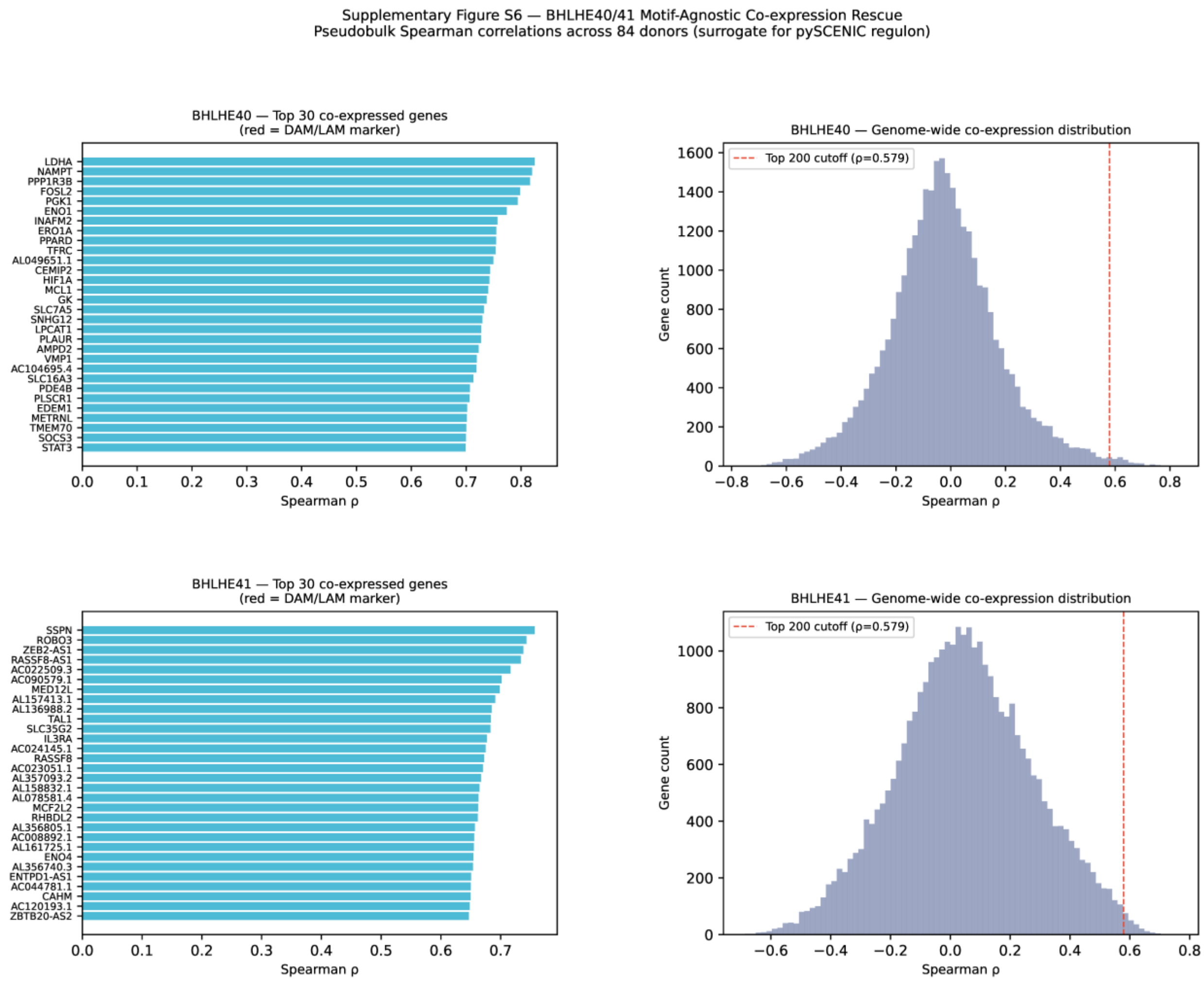


***Supplementary Figure S2. BHLHE40/41 motif-agnostic pseudobulk co-expression analysis.***

(A, C) Top 30 genes positively co-expressed with BHLHE40 and BHLHE41, respectively, across 84 donor pseudobulk samples (Spearman ρ); bars coloured to indicate whether the gene is a canonical DAM/LAM marker (TREM2, SPP1, CD9, GPNMB, LGALS3, LPL, APOE, CCL3L1, CCL4L2, RGCC). All 30 bars share the same colour for both TFs, confirming the absence of DAM/LAM markers among the top co-expressed genes. (B, D) Genome-wide Spearman correlation distributions for BHLHE40 and BHLHE41 respectively; distributions are symmetric and centred near zero; dashed vertical line indicates the top-200 cutoff ($\rho \approx 0.579$ for both TFs). Fisher's exact test found no significant enrichment of DAM/LAM marker genes within the top 200 co-expressed genes for either TF (BHLHE40: OR = 0.00, P value = 1.0; BHLHE41: OR = 0.00, P value = 1.0), indicating that BHLHE40/41 donor-level expression variance is not coupled to the DAM/LAM transcriptional programme across individuals.

The null result is consistent with BHLHE40/41 operating within cell-state transitions (captured by cell-level analyses) rather than as donor-level expression axes (captured by the pseudobulk approach). The genome-wide correlation profiles and top co-expressed gene lists are provided for exploratory reference; this analysis constitutes a surrogate regulon approximation and does not substitute for a cisTarget-validated pySCENIC regulon.

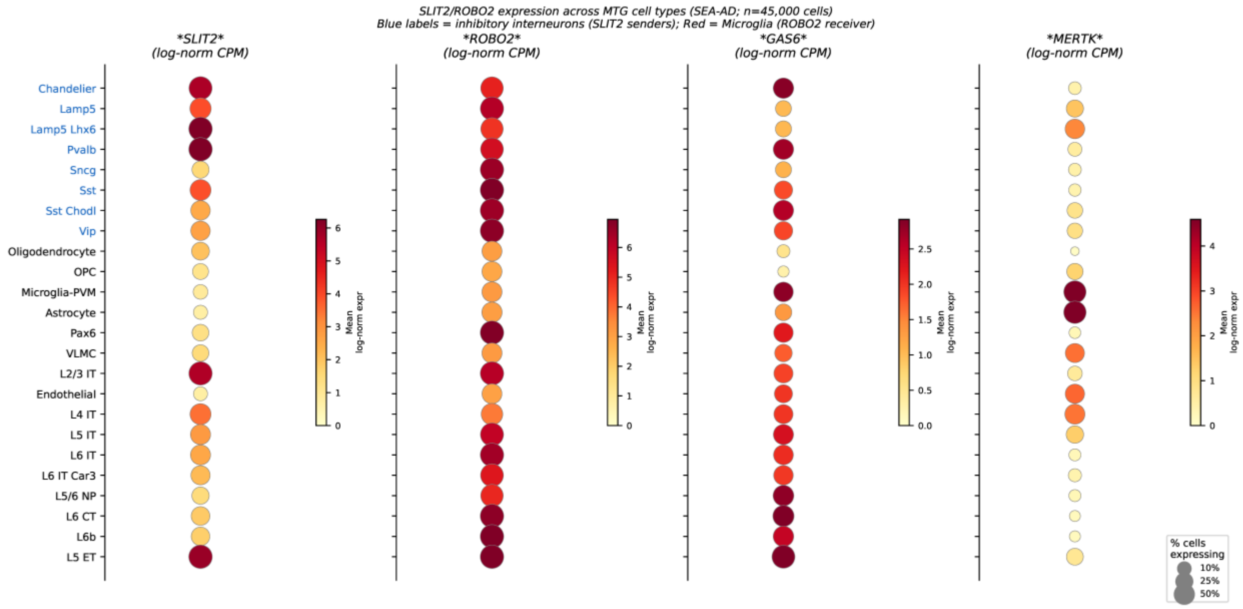


***Supplementary Figure S3. SLIT2 and ROBO2 expression across MTG cell types.***

Dot plot showing log-normalised expression of SLIT2, ROBO2, GAS6, and MERTK across MTG cell types in the SEA-AD MTG (n = 45,000 nuclei subsampled; 5,000 cells per broad CellChat cell type; plotted at fine-grained cell type resolution). Dot diameter encodes the percentage of cells expressing the gene (% cells with > 0 counts); dot colour encodes mean log-normalised expression (yellow–orange–red scale, 0–95th percentile). All cell type labels are in black. A dashed horizontal line separates the inhibitory interneuron subtypes (above the line; SLIT2/GAS6 senders, labelled with a bracket annotation “Inhibitory interneurons”) from other cell types (below the line, including Microglia-PVM). Cell types are ordered with inhibitory interneurons at the top, followed by non-neuronal and glial types (including Microglia-PVM), then excitatory neuron subtypes at the bottom. SLIT2 shows highest expression in Chandelier, Lamp5 Lhx6, Pvalb, and Sst interneurons. ROBO2 is broadly expressed with the highest percentage of expressing cells in Microglia-PVM. MERTK expression is concentrated in Microglia-PVM, consistent with its role as a microglial efferocytosis receptor. This figure provides expression-level support for the SLIT2→ROBO2 and GAS6→MERTK L-R interactions identified by CellChat; it does not constitute functional validation of these interactions.

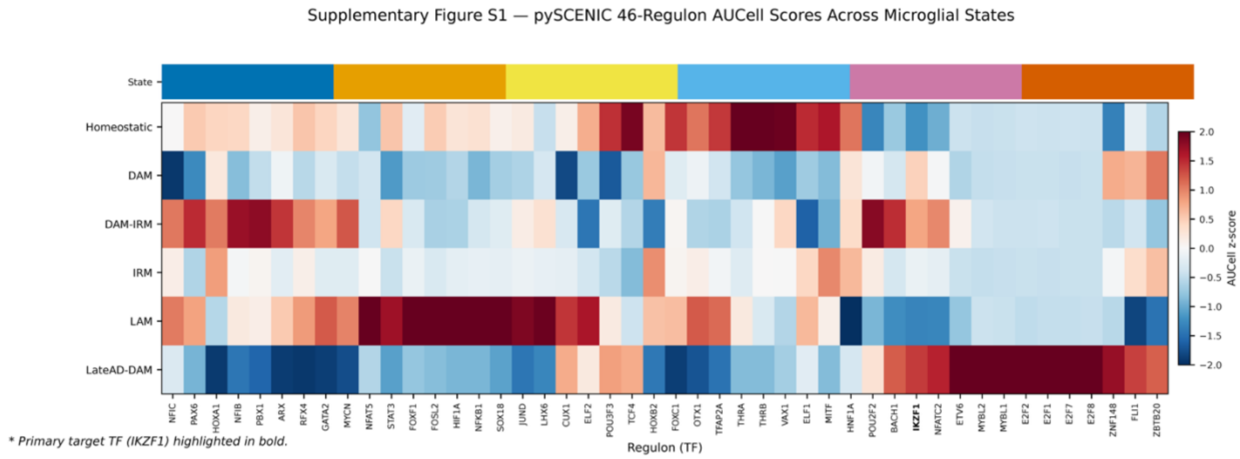


***Supplementary Figure S4. pySCENIC regulon activity heatmap across microglial substates.***

Heatmap of scaled AUCell regulon activity scores for all 46 RcisTarget-pruned pySCENIC regulons (columns) across six microglial substates (rows). Values are mean scaled AUCell scores per substate. Diverging colour scale: blue = below-mean activity, red = above-mean activity; vertical colourbar on the right. A narrow colour strip immediately above the heatmap rows uses distinct colours to indicate substate identity (matching the y-axis row labels). IKZF1(+) is visible near the right-hand end of the x-axis and shows elevated activity in LateAD-DAM. The IKZF1(+) regulon is marked with an asterisk (*) in the x-axis label; all other labels are in black. The heatmap illustrates the breadth of the 46-regulon landscape and the state-specificity of IKZF1 relative to other regulons.

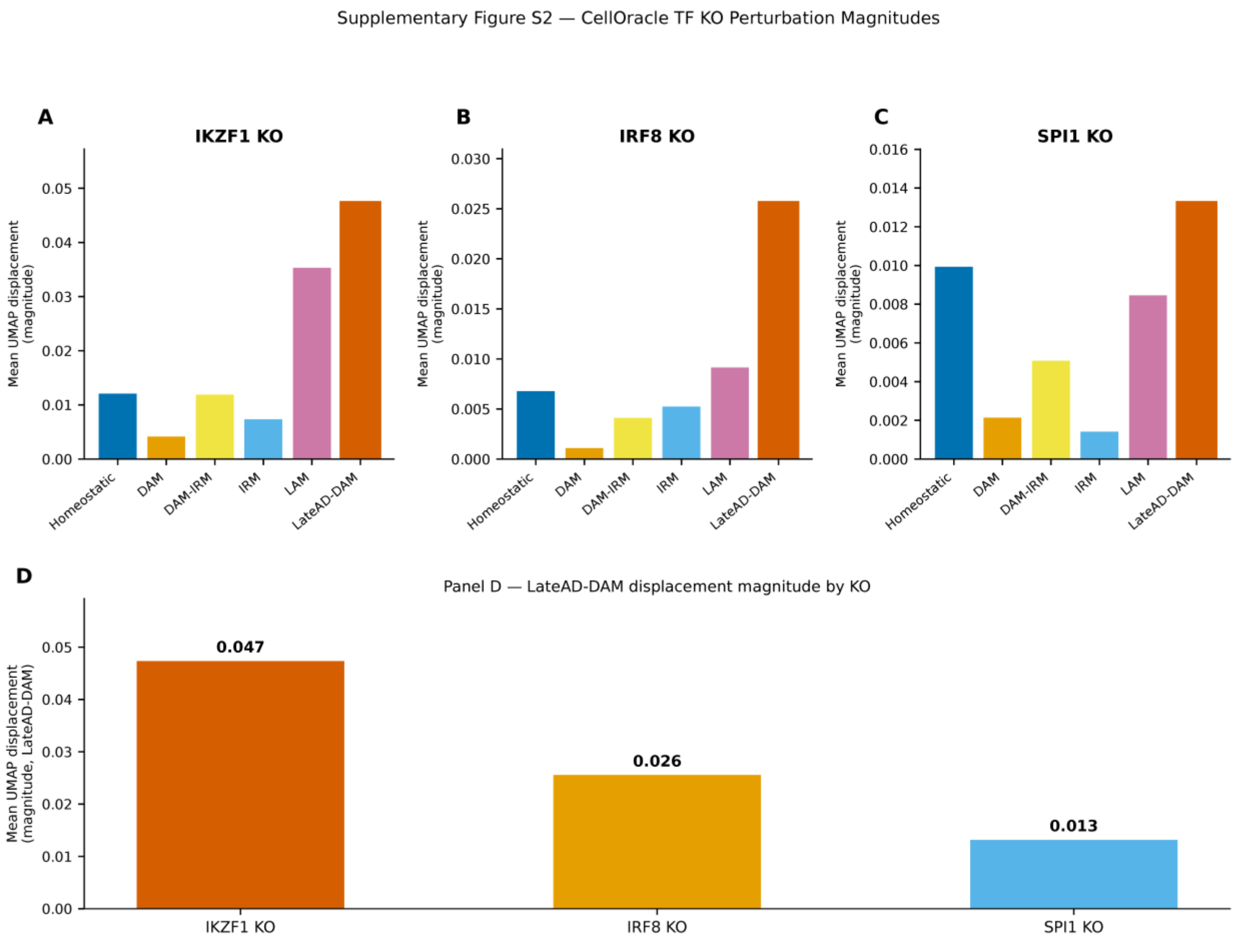


***Supplementary Figure S5. CellOracle in-silico TF knockout perturbation magnitudes across microglial substates.***

Bar charts of mean UMAP displacement magnitude (|ΔUMAP| per cell) across six microglial substates after in-silico knockout of (A) IKZF1, (B) IRF8, and (C) SPI1; bars colour-coded and hatch-coded by substate. (D) Direct comparison of LateAD-DAM 2D UMAP displacement magnitude across the three KO conditions; IKZF1 KO ranks highest (0.047), followed by IRF8 KO (0.026) and SPI1 KO (0.013). Values are in UMAP units and represent an ordinal ranking within this embedding — they should not be interpreted as absolute effect sizes, as UMAP is not metric-preserving. Analyses were performed on a 20,000-cell stratified subsample (≥ 200 cells per state; random_state = 42). The GRN used for CellOracle was pruned to the 100K-cell pySCENIC consensus network.

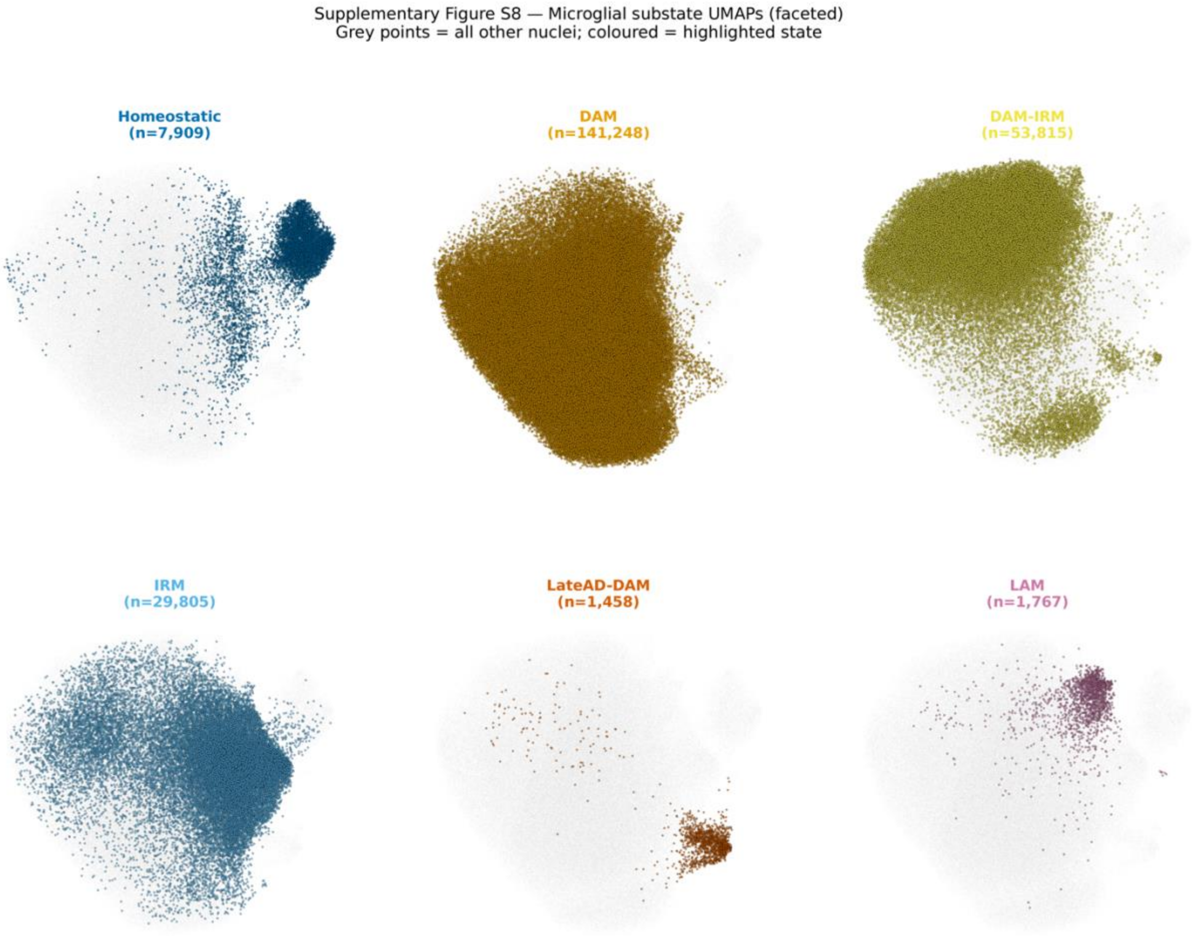


***Supplementary Figure S6. Microglial substate UMAPs (faceted).***

Small-multiple UMAP showing each of the six microglial substates individually (2 × 3 grid). In each panel, nuclei belonging to the highlighted substate are shown in the substate colour with thin semi-transparent borders; all remaining nuclei are grey. Cell counts per substate are shown in each panel title. This figure is an accessibility companion to Figure 1A, allowing readers to assess the UMAP position of each substate without relying on colour differentiation across overlapping clusters.

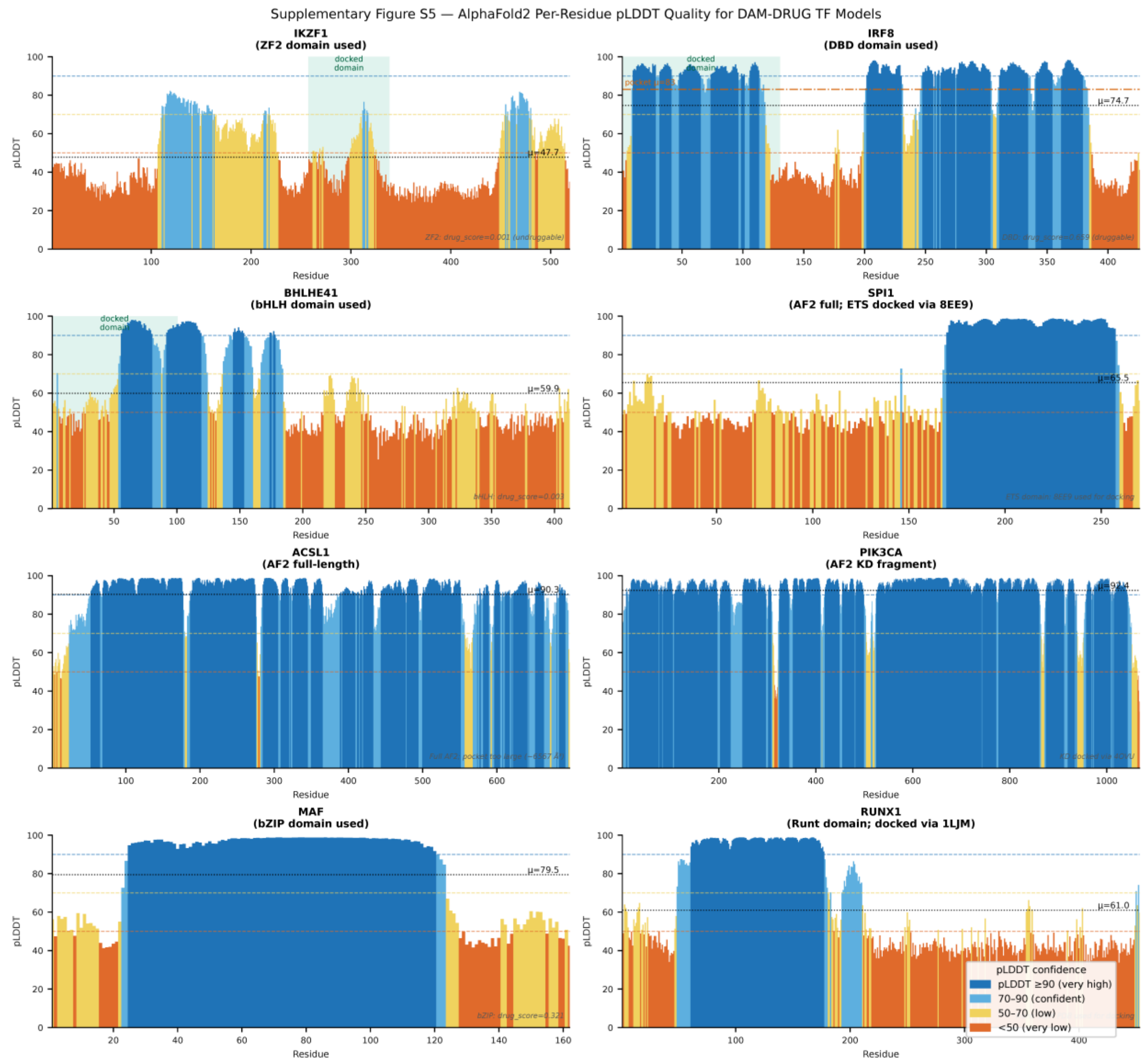


***Supplementary Figure S7. AlphaFold2 per-residue pLDDT quality scores for all DAM-DRUG target TF structural models.***

Per-residue pLDDT confidence scores for the structural models used in the virtual screening pipeline.

AlphaFold2 models (no experimental structure used for docking): IKZF1 ZF2 domain (mean pLDDT = 41.7; orthosteric docking not pursued due to negligible druggability — drug_score = 0.001), IRF8 DBD (mean = 74.1; no experimental human IRF8 DBD structure in the PDB; benchmarked against IRF1 PDB 1IF1, TM-score = 0.90), BHLHE41 bHLH domain (mean ≈ 39.6; deprioritised by fpocket drug_score = 0.003), ACSL1 full-length AF2 model (shown for reference; not taken to docking), PIK3CA kinase domain AF2 fragment (shown for reference; not taken to docking).

Experimental PDB structures (docking used experimental coordinates): RUNX1 Runt domain (PDB 1LJM, 2.7 Å resolution; pLDDT profile shown for comparison only), SPI1

ETS domain (PDB 8EE9; pLDDT profile shown for comparison only), MAF bZIP domain (homology model based on PDB 4EOT; mean pLDDT = 79.5; drug_score = 0.432).

PPARG is not shown in this figure because docking used the experimental structure PDB 1FM9 exclusively and no AF2 model was assessed for this target.

Colour coding: blue = pLDDT ≥ 90 (very high confidence), light blue = 70–90 (confident), yellow = 50–70 (low), orange < 50 (very low). Shaded regions indicate the specific domain used for pocket detection and docking. Dashed horizontal lines mark the per-protein mean pLDDT. Low mean pLDDT for IKZF1 (41.7) and BHLHE41 (39.6) is consistent with their intrinsically disordered character and explains the negligible fpocket druggability scores for these targets.

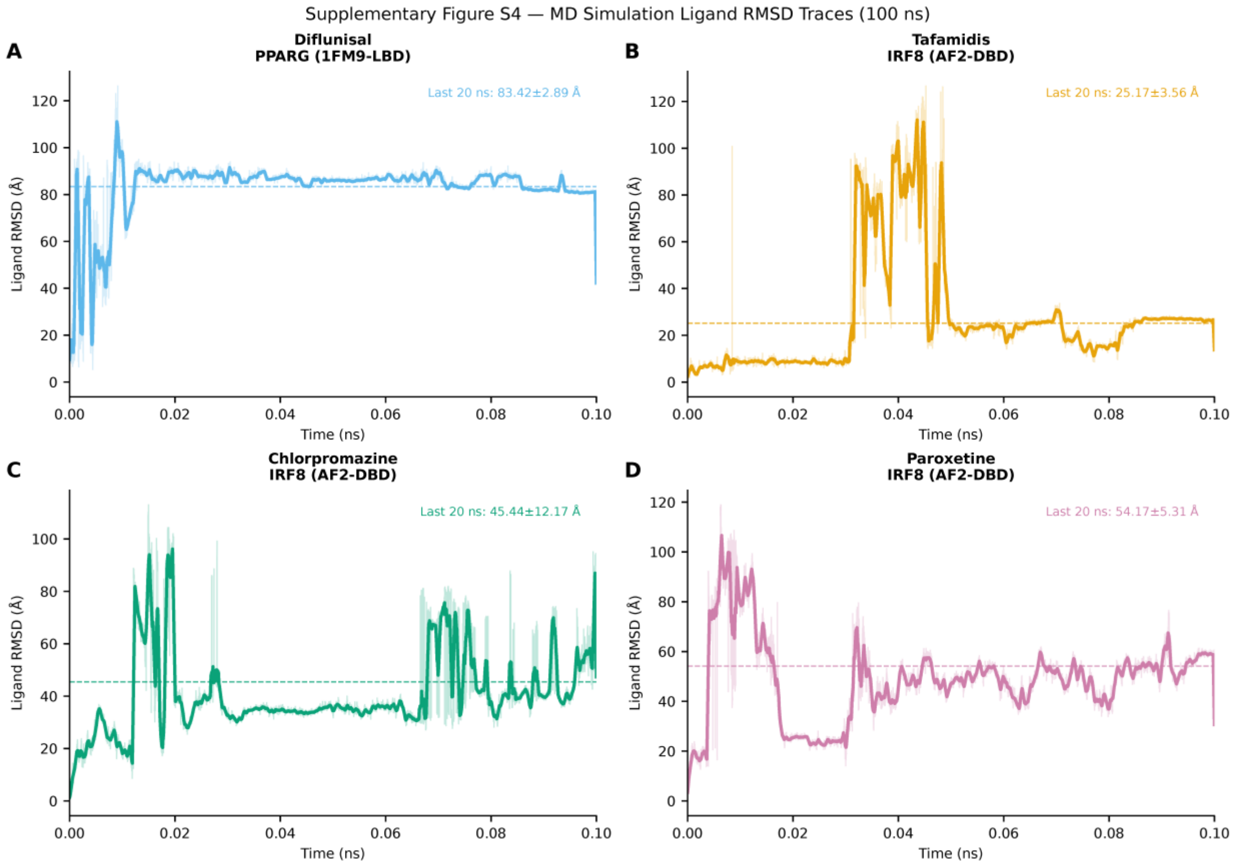


***Supplementary Figure S8. Ligand RMSD traces from 100 ns MD simulations for top shortlisted compounds.***

Ligand RMSD (Å) relative to the initial docked pose over 100 ns MD simulation for (A) diflunisal bound to PPARG LBD (experimental structure PDB 1FM9, 2.1 Å resolution; last 20 ns mean = 83.42 ± 2.89 Å), (B) tafamidis bound to IRF8 DBD (AlphaFold2 model; no experimental human IRF8 DBD structure available; last 20 ns mean = 25.17 ± 3.56 Å), (C) chlorpromazine bound to IRF8 DBD (AlphaFold2 model; last 20 ns mean = 45.44 ± 12.17 Å), and (D) paroxetine bound to IRF8 DBD (AlphaFold2 model; last 20 ns mean

= 54.17 ± 5.35 Å). Dashed horizontal line indicates the last-20-ns mean; shaded band = ± 1 SD. All four compounds show ligand RMSD substantially exceeding the 3–4 Å stability threshold, indicating departure from the initial docked pose. These results are interpreted as a collective limitation of the implicit-solvent MM-GBSA pipeline rather than as evidence of stable binding; all four compounds are classified as low-confidence computational hypotheses requiring experimental triage (SPR/TSA).

MD simulations were performed with GROMACS 2024 (AMBER14 force field + GBn2 implicit solvent; 200-step energy minimisation prior to production run; 1 ns equilibration; 100 ns production; 250 frames sampled). Ligand RMSD was computed relative to frame 0 of the production trajectory after backbone alignment to the protein core (GROMACS rms module).
